# Phylogeny-corrected identification of microbial gene families relevant to human gut colonization

**DOI:** 10.1101/189795

**Authors:** Patrick H. Bradley, Stephen Nayfach, Katherine S. Pollard

## Abstract

The mechanisms by which different microbes colonize the healthy human gut versus other body sites, the gut in disease states, or other environments remain largely unknown. Identifying microbial genes influencing fitness in the gut could lead to new ways to engineer probiotics or disrupt pathogenesis. We approach this problem by measuring the statistical association between having a species having a gene and the probability that the species is present in the gut microbiome. The challenge is that closely related species tend to be jointly present or absent in the microbiome and also share many genes, only a subset of which are involved in gut adaptation. We show that this phylogenetic correlation indeed leads to many false discoveries and propose phylogenetic linear regression as a powerful solution. To apply this method across the bacterial tree of life, where most species have not been experimentally phenotyped, we use metagenomes from hundreds of people to quantify each species’ prevalence in and specificity for the gut microbiome. This analysis reveals thousands of genes potentially involved in adaptation to the gut across species, including many novel candidates as well as processes known to contribute to fitness of gut bacteria, such as acid tolerance in Bacteroidetes and sporulation in Firmicutes. We also find microbial genes associated with a preference for the gut over other body sites, which are significantly enriched for genes linked to fitness in an in vivo competition experiment. Finally, we identify gene families associated with higher prevalence in patients with Crohn’s disease, including Proteobacterial genes involved in conjugation and fimbria regulation, processes previously linked to inflammation. These gene targets may represent new avenues for modulating host colonization and disease. Our strategy of combining metagenomics with phylogenetic modeling is general and can be used to identify genes associated with adaptation to any environment.

**Author Summary:** Why do certain microbes and not others colonize our gut, and why do they differ between healthy and sick people? One explanation is the genes in their genomes. If we can find microbial genes involved in gut adaptation, we may be able to keep out pathogens and encourage the growth of beneficial microbes. One could look for genes that were present more often in prevalent microbes, and less often in rare ones.

However, this ignores that related species are more likely to share an environment and also share many unrelated phenotypes simply because of common ancestry. To solve this problem, we used a method from ecology that accounts for phylogenetic relatedness. We first calculated gut prevalence for thousands of species using a compendium of shotgun sequencing data, then tested for genes associated with prevalence, adjusting for phylogenetic relationships. We found genes that are associated with overall gut prevalence, with a preference for the gut over other body sites, and with the gut in Crohn’s disease versus health. Many of these findings have biological plausibility based on existing literature. We also showed agreement with the results of a previously published high-throughput screen of bacterial gene knockouts in mice. These results, and this type of analysis, may eventually lead to new strategies for maintaining gut health.

## 1 Introduction

Microbes that colonize the human gastrointestinal (GI) tract have a wide variety of effects on their hosts, ranging from beneficial to harmful. Increasing evidence shows that commensal gut microbes are responsible for training and modulating the immune system [1, 2], protecting against inflammation [3] and pathogen invasion (reviewed in Sassone-Corsi and Raffatellu [4]), affecting GI motility [5], maintaining the intestinal barrier [6], and potentially even affecting mood [7]. In contrast, pathogens (and conditionally-pathogenic microbes, or “pathobionts”) can induce and worsen inflammation [8, 9], increase the risk of cancer in mouse models [10], and cause potentially life-threatening infections [11]. Additionally, the transplantation of microbes from a healthy host (fecal microbiota transplant, or FMT) is also a highly effective therapy for some gut infections [12], although it is still an active area of investigation why certain microbes from the donor persist long-term and others do not [13], and how pre-existing inflammatory disease affects FMT efficacy [14]. Which microbes are able to persist in the GI tract, and why some persist instead of others, is therefore a question with consequences that directly impact human health.

Because of this, we are interested in the specific mechanisms by which microbes colonize the gut, avoiding other potential fates such as being killed in the harsh stomach environment, simply passing through the GI tract transiently, or being outcompeted by other gut microbes. Understanding these mechanisms could yield opportunities to design better probiotics and to prevent invasion of the gut community by pathogens. In particular, creating new therapies, whether those are drugs, engineered bacterial strains, or rationally designed communities, will likely require an understanding of gut colonization at the level of individual microbial genes. We also anticipate that these mechanisms may vary in health vs. disease, since, for example, different selective pressures are known to be present in inflamed versus healthy guts [15, 16].

One approach that has been used to link genetic features to a phenotype is to correlate the two using observational data. Most typically, this approach is applied in the form of genome-wide association mapping, in which phenotypes are correlated with genetic markers across individuals in a population. While we are interested in comparing phenotypes and genetic features *across*, rather than within species, the approach we take in this paper is conceptually similar. In order to perform association mapping, it is necessary to account for population structure, that is, dependencies resulting from common ancestry; otherwise, spurious discoveries can be made in genome-wide association studies [17]. Analogously, we expected it to be important to choose a method that can account for the confounding effect of phylogeny when testing for associations across species.

There is increasing interest in using phylogenetic information to make better inferences about associations between microbes and quantities of interest. For example, co-conservation patterns of genes (“correlogs”) have been used to assign functions to microbial genes [18], and genome-wide association studies have been applied within a genus of soil bacteria [19] as well as across strains of *Neisseria meningitidis* [20]. Recent publications have also described techniques that use information from the taxonomic tree to more accurately link clades in compositional taxonomic data to covariates [21–23]. However, so far, only one study has attempted to associate genes with a preference for the gut [24]. That study introduced a valuable method based on UniFrac and gene-count distances, which compares how well gut- vs. non-gut-associated microbes cluster on the species tree compared to a composite gene tree. This study also provides an important insight in the form of evidence of convergence of glycoside hydrolase and glycosyltransferase repertoires among gut bacteria, suggesting horizontal gene transfer within the gut community to deal with a common evolutionary pressure. The method described in that study, though, requires a binary phenotype of gut presence vs. absence. Deciding which microbes are “gut” vs. “non-gut” requires manual curation and can be somewhat subjective, as microbes have a continuous range of prevalences and can appear in multiple environments; this binarization could also potentially decrease power by excluding microbes with intermediate phenotypes. The method also requires multiple sequence alignments and trees to be built for every gene family under analysis, which are computationally intensive to generate over a large set of genomes.

We take a complementary approach and use a flexible technique, known as phylogenetic linear modeling, to detect associations between microbial genotype and phenotype while accounting for the fact that microbes are related to one another by vertical descent. Phylogenetic linear models have an extensive history in the ecology literature dating back to seminal works by Felsenstein [25] and Grafen [26]. However, despite their power, genome-scale applications of these models are still few in number [27] and, with the exception of one recent study that applied phylogenetic linear modeling to newly-sequenced isolate genomes from plant-associated microbial communities [28], have typically been used to relate traits of macroorganisms (e.g., anole lizards [29]) to their genotypes. While there is a growing appreciation for the need to explicitly account for phylogeny in microbial community analyses [27, 30], we believe ours is the first study to directly apply this class of methods to metagenomic data.

This approach to accounting for phylogenetic relationships is general and could be applied to measure association of any quantitative phenotype with genotypes or other binary or quantitative characteristics. In this study, we focus on phenotypes related to the ability of bacteria to colonize the human gut: 1. overall prevalence in the guts of hosts from a specific population (e.g., post-industrialized countries), which we expect to capture ease of transmission, how cosmopolitan microbes are, and how efficiently they colonize the gut; 2. a preference for the gut over other human body sites in the same hosts, which we expect to capture gut colonization more specifically; and 3. a preference for the gut in disease (e.g., Crohn’s disease) versus health. We present a novel analytic pipeline in which we estimate these quantitative phenotypes for thousands of bacterial species directly from existing shotgun metagenomics data, both obviating the need for us to draw a cutoff between “gut” and “non-gut” microbes, and also giving us the necessary power to detect associations (Fig S2). Coupling these phenotype estimates with phylogenetic linear models, we generate a compendium of thousands of bacterial genes whose functions may be involved in colonizing the human gut.

## 2 Results

We present a phylogeny-aware method for modeling associations between the presence of specific genes in bacterial genomes and quantitative phenotypes that measure how common these species are in the human microbiome. To apply phylogenetic linear modeling to the microbiome, we needed to solve three problems. First, we had to show that these models controlled false positives and had reasonable power on large bacterial phylogenies. Second, we needed to develop estimators that captured meaningful phenotypes related to bacterial colonization of humans for thousands of diverse bacterial species, most of which have never been studied in isolation, much less experimentally assayed for their abilities to colonize a mammalian body site. The third problem was to estimate genotypes (e.g., gene presence-absence) for each species. The analysis framework we describe is quite general and could be easily extended to link other phenotypes to genotypes across the tree of life.

### 2.1 Phylogenetic linear models solve the problem of high false positive rates when testing for associations on bacterial phylogenies

To test for associations between quantitative phenotypes and binary genotypes across species, we use models with the following form:

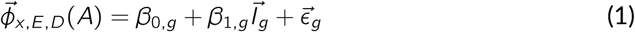

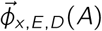 is a vector of quantitative phenotypes of interest, assessed in one environment *e*_*x*_ out of a set of possible environments *E*, normalizing out a set of study effects *D*, estimated from the dataset *A*. The elements of the vector 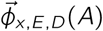 are *ϕ*_*m,x,E,D*_(*A*), the phenotype value for microbe *m. β*_0,*g*_ is a baseline phenotype value, *β*_1,*g*_ is the effect of gene *g* on 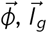 is a vector whose elements *I*_*m,g*_ are 0 if gene *g* is absent in species *m* and 1 if present, and 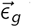 is the remaining unmodeled variation in 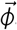. We fit one model per gene *g*. The distribution of the residuals 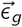 is the key difference between standard and phylogenetic linear models. In the standard model, the residuals are assumed to be independent and normally distributed. In the phylogenetic model, however, the residuals covary, with more closely-related species having greater covariance (see Methods, 4.5 and 4.6; for a glossary of notation, see Methods, 4.17).

To explore the potential pitfalls of failing to correct for phylogenetic structure in cross-species association tests, we generated a species tree for thousands of bacteria with genome sequences (see Methods, 4.3). In order to have a consistent operational definition of a microbial species, we used a set of previously defined bacterial taxonomic units with approximately 95% pairwise average nucleotide identity across the entire genome [31]. The methods we describe can be applied to other taxonomic levels or with other species definitions. Using this species tree, we performed simulations (see Methods, 4.11) for each of the four major bacterial phyla in the human gut (Bacteroidetes, Firmicutes, Proteobacteria, and Actinobacteria [32]). Specifically, we generated simulated phenotypes along the species tree, and then, for each phenotype, simulated a binary genotype for each species that covaried with the phenotype to varying degrees, including no association. We used levels of covariation spanning those we observed empirically between prevalence of species in gut metagenomes and presence-absence of genes (see below). (An effect size of 0.5 corresponds approximately to a 50% increase in prevalence, while an effect size of 1.0 corresponds approximately to a 100% increase in prevalence: see Fig S8.) These binary genotypes also had varying levels of overall phylogenetic signal (Ives-Garland *α*).

We then fit phylogenetic and standard linear models to the simulated data and tested for a relationship between each binary genotype and its corresponding continuous phenotype. For both standard and phylogenetic linear models, separate models were fit for each of the four phyla. The results were used to estimate false positive rate (Type I error) and power (1 - Type II error) for the two methods across different effect sizes.

These analyses showed that standard linear models result in many false positive associations. When the binary genotype was specified to be wholly uncorrelated (i.e., under the null), *p*-values from the linear model showed a strong anticonservative bias (Fig 1B, D) with many more significant *p*-values than expected under no correlation. While lower levels of phylogenetic signal (larger Ives-Garland *α*) did result in less bias in the standard linear model, the false positive rate remained over 25% at *p* = 0.05. In contrast, the phylogenetic linear model *p*-value distribution was flat and Type I error was controlled appropriately (Fig 1A, C). This means that at the same *p*-value threshold, linear models will identify many spurious relationships compared to phylogenetic linear models. Further, our simulations with non-zero associations showed that the phylogenetic model has high power when applied to gut bacterial phyla, even for small effect sizes (Fig 1E; see Methods 4.11). These results emphasize the importance of using models that account for phylogenetic relationships in cross-species association testing and demonstrate the feasibility of applying phylogenetic linear models to the human microbiome.

**Figure 1.**
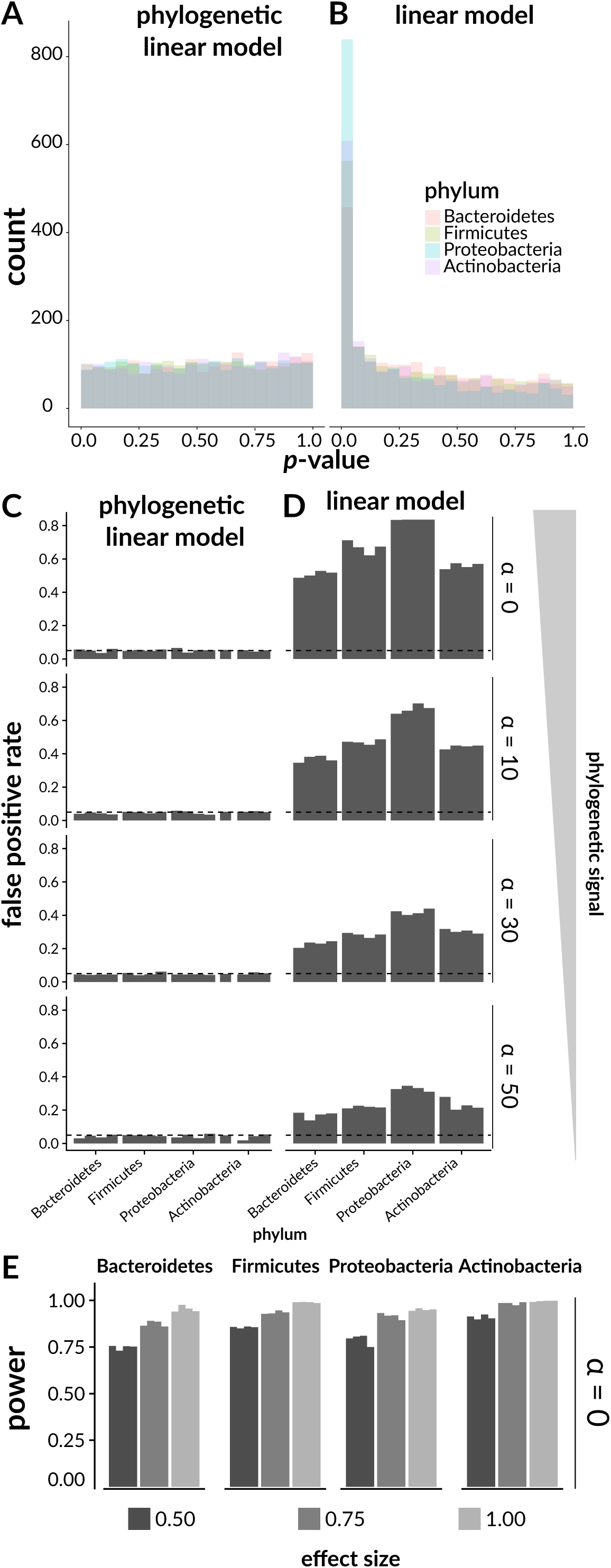
Failing to account for tree structure results in an elevated false positive rate. Continuous phenotypes and binary genotypes were simulated across the trees for the four phyla under consideration. A-B show results for the null of no true phenotype-genotype correlation. A) Histogram of *p*-values for simulated phenotypes and genotypes on the Bacteroidetes tree, using phylogenetic (left) or standard (right) linear models. The phylogenetic model distribution was similar to a uniform distribution, while the standard model was very anticonservative, having an excess of small *p*-values. B) False positive rate (Type I error rate) at *p* = 0.05 for the phylogenetic and standard models. C) Traits with varying levels of “true” association spanning values we observed in real data were simulated, and power was computed using phylogenetic linear models.

### 2.2 Estimating quantitative phenotypes from shotgun data

To apply phylogenetic linear modeling to the microbiome we sought to define meaningful phenotypes for thousands of bacterial species, all of which have genome sequences but most of which have never been experimentally tested for, e.g., their abilities to grow on particular substrates or to colonize a model mammalian gut. We hypothesized that the prevalence and specificity of bacterial species in an environment, such as the human gut, should relate to their ability to colonize that environment and to how well adapted they are to persist there. These quantities can be thought of as phenotypes that can be estimated directly from shotgun metagenomics data. The precise taxonomic composition of a healthy gut microbiome can vary significantly from person to person [33], indicating that the ability of a microbe to colonize the gut is quantitative (and likely context-dependent, and stochastic). This phentoype can be conceptualized differently depending on which aspects of colonization one wishes to capture. We present metagenome-based estimators for two different types of colonization phenotpyes. These are described in the context of our goal of studying the gut microbiome, but the approach is general and could be used to quantify how well a given genotype discriminates species found in or specific to any environment.

The first phenotype is the probability of observing a microbial species *m* in an environment *e*_*x*_, that is, its *overall prevalence P* (*m* | *e*_*x*_). Both genes relating to survival in the GI tract and genes relating to survival, persistence, and dispersal in the outside environment are expected to correlate with overall prevalence. Prevalence can be estimated by the frequency with which the species is observed in a sample from the environment, for example, using a logit transform to enable linear modeling and pseudocounts to avoid estimates of 0 or 1 (see Methods, 4.8).

The second type of quantitative phenotype is the *environmental specificity* of a microbial species, which we define as the conditional probability that a sample is derived from one environment in a set of environments, given that the species is present in the sample. This parameter captures the power of a given microbe as a marker to discriminate between two or more different environments, such as different body sites or types of hosts (see Methods, 4.9). This is distinct from its overall prevalence in the environment.

We developed an estimator for environmental specificity and applied it to two separate gut microbial phenotypes. First, we considered a phenotype defined as the conditional probability that a given body site is the gut and not another body site, given that a particular species is present. The physical distance between body sites is much smaller than the distance between hosts, and microbes from one body site are likely to be transiently introduced to others. Hence, enrichment of a species in one body site over others is stronger evidence for selection (versus dispersal) than is overall prevalence in that body site alone. We estimate this parameter with a *body-site specificity score* that uses metagenomics data to measure how predictive a particular microbe is for the gut versus other body sites (e.g., skin, urogenital tract, oropharynx, or lung).

The second type of environmental specificity we considered is the conditional probability that a host has a disease given that a particular species is present. This *disease-specific specificity score* is estimated in a similar way to the body-site specificity score (see Methods, 4.9). We focus on Crohn’s disease, a type of inflammatory bowel disease known to be associated with dramatic shifts in the gut microbiota and in gut-immune interactions [34]. Genes associated with this disease-specific prevalence could illuminate differences in selective pressures between healthy vs. diseased gut environments. Both scores are based on maximum *a posteriori* (MAP) estimates of the conditional probability of a sample being from the gut given that a microbe is observed in the sample. To account for sampling noise, we use a shrunken estimate with a Laplace prior (see Methods, 4.9).

### 2.3 Genes associated with species prevalence in healthy human gut metagenomes

We assembled a compendium of published DNA sequencing data from healthy human stool microbiomes across five studies in North America, Europe, and China (433 subjects total). Using the MIDAS database and pipeline [31], we mapped metagenomic sequencing reads from each run to a panel of phylogenetic marker genes, and from these, estimated species relative abundances. Multiple runs corresponding to the same individual were averaged. We then estimated the prevalence (probability of non-zero abundance) of each species across these subjects, weighting each study equally and adding pseudocounts to avoid probabilities of exactly 0 or 1 (see Methods, 4.8). Finally, we determined whether genes (here, we take “genes” to mean members of a FIGfam protein family, which are designed to approximate “isofunctional homologs” [35]) were present or absent in the pangenomes of each species, based on sequenced genomes included in the MIDAS database, such that any FIGfam annotated in at least one sequenced isolate was considered to be present in the pangenome. This approach to genotyping could be extended to additionally include single-amplified genomes and metagenome-assembled genomes (see Discussion). Our analysis framework can also be applied to genotypes other than gene presence-absence (e.g., nucleotide or amino acid changes). We note that while the FIGfam database does include many hypothetical protein families of unknown function, many bacterial genes lack even this level of annotation, so a more comprehensive grouping of genes into orthologous or functionally homologous groups could reveal yet more novel associations.

As expected, the most prevalent species overall included *Bacteroides vulgatus, Bacteroides ovatus*, and *Faecalibacterium prausnitzii*, while the least prevalent included halophiles and thermophiles (Table S1). Gut prevalence had a strong phylogenetic signal (Pagel’s *λ* = 0.97, likelihood-ratio *p <* 10^−22^), meaning that it was strongly correlated with the evolutionary relatedness of species. This emphasizes the need for phylogeny-aware modeling so that signal linking genes to prevalence will not be drowned out by shared variation in gene content between closely-related species.

To demonstrate the effect of phylogenetic correlation empirically, we fit both a standard linear model and a phylogenetic linear model for each of the four common gut phyla and all genes present in that phylum. These models relate logit-transformed estimates of the prevalence of different species in a phylum to a gene’s presence-absence in those species’ pangenomes. Recall that the residual variation in logit-prevalence is independent and normally distributed in the standard linear model, but has a distribution encoding correlations proportional to species relatedness in the phylogenetic linear model (see Methods, 4.6). For both standard and phylogenetic linear models, separate models were fit for each phylum. While this means that genes weakly-associated across the entire tree of life may have been missed by this approach, it has the advantage of both reducing the memory needed to store the gene presence-absence matrix and allowing for phylum-specific rates of evolution for our phenotype of interest. We modeled associations for 144,651 genes total across the four phyla, fitting 381,846 models total (since some genes are present in multiple phyla).

We used the parameter estimates and their standard errors from fitted models to test null hypotheses of the form *H*_0_ : *β*_1,*g*_ = 0, meaning gene *g* is not associated with gut prevalence of species in a particular phylum. The *p*-values were adjusted for multiple testing using the false discovery rate (FDR) (see Methods, 4.6). We found 9,830 FIGfam gene families positively associated with logit-prevalence within at least one phylum (FDR *q* ≤ 0.05) using phylogenetic linear models, 47% of which had no annotated function. We observed that 75% of the significant genes from these tests had effect sizes larger than (Bacteroidetes) 0.93, (Firmicutes) 1.03, (Proteobacteria) 0.35, and (Actinobacteria) 2.04, which are within the range of effect sizes for which phylogenetic linear models showed good performance in simulations (see above).

With standard linear models our tests identified 25,185 genes associated with gut prevalence, substantially more than with phylogenetic linear models (17.4% versus 6.8% of total). Based on our simulations, these likely included many false positives. The top results of phylogenetic versus standard linear models (Fig 2) illustrate the pitfalls of not correcting for phylogenetic correlation. Using the standard model, we recover associations such as those seen in Fig 2A-B: a subunit of dihydroorotate dehydrogenase in Bacteroidetes (Fig 2B) and in Firmicutes, a particular type of glutamine synthetase (Fig 2A). While these associations might look reasonable at a first glance, on closer inspection, they depend on the fact that these genes are near-uniformly present in entire clades of bacteria. These clades are, in general, more prevalent in the gut compared to the rest of the species in the tree. However, any finer structure relating to differences between close neighbors is lost. For example, dihydroorotate dehydrogenase (Fig 2B) is found not only in the human gut commensal *Bacteroides caccae*, but also in its relative *Bacteroides reticulotermitis*, which was not only low-prevalence in our samples but was indeed isolated from the gut of a subterranean termite [36].

**Figure 2.**
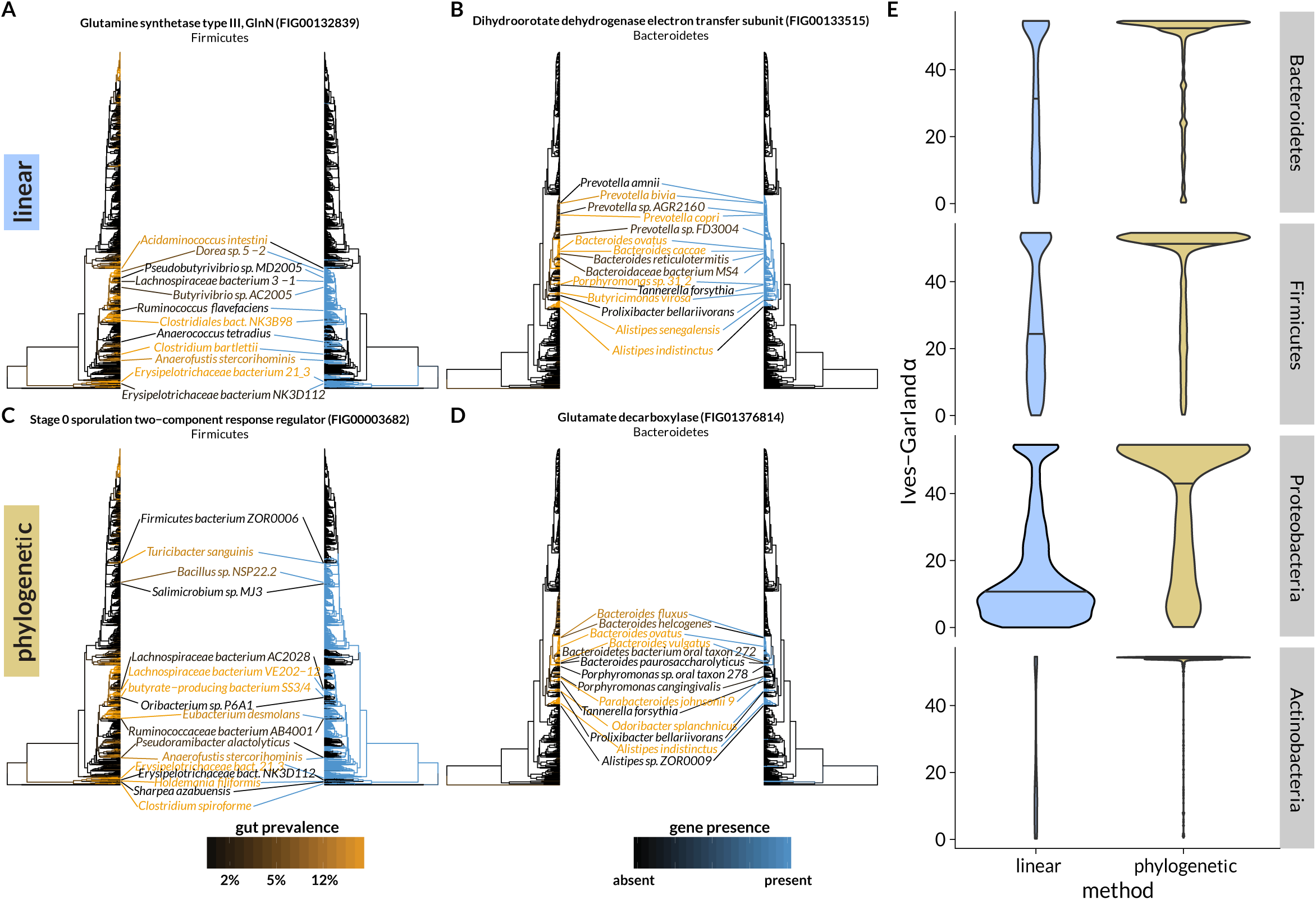
Examples of hits from standard linear (blue highlights) and phylogenetic (orange highlights) models. In each panel, the tree on the left is colored by species prevalence (black to orange), while the tree on the right is colored by gene presence-absence (blue to black). Selected species are displayed in the middle; lines link species with the leaves to which they refer. The color of the line matches the color of the leaf. A-B) The standard model recovered hits that matched large clades but without recapitulating fine structure. C-D) The phylogenetic model recovered associations for which more of the fine structure was mirrored between the left-hand and right-hand trees, as exemplified by the species labeled in the middle. E) Violin plots of Ives-Garland *α*, a summary of the rate of gain and loss of a binary trait across a tree, for genes significantly associated with prevalence in the standard (left, blue) and phylogenetic (right, orange) linear models. Horizontal lines mark the median of the distributions. The phylogenetic (orange) and standard linear (blue) models were significantly different for each phylum (Wilcox test for Bacteroidetes: 4 *×* 10^−6^; Firmicutes: 7 *×* 10^−11^; Proteobacteria: 2 *×* 10^−22^; Actinobacteria: 2 *×* 10^−22^).

While this alone does not necessarily constitute evidence *against* these genes having adaptive functions in the human gut, we do expect that matched pheno- and genotypic differences between close phylogenetic neighbors offer stronger evidence for an association. An analogy can be drawn with genome-wide association mapping in humans: models that do not account for correlations between sites caused by population structure, as opposed to selective pressure, will tend to identify more spurious associations. In contrast, because the phylogenetic null model “expects” phenotypic correlations to scale with the evolutionary distance between species, this approach will tend to upweight cases where phylogenetically close relatives have different phenotypes and where distant relatives have similar phenotypes. This leads to the identification of candidate genes that capture more variation between close neighbors (Fig 2C-D). Thus, phylogenetic linear models will identify genes whose presence in genomes is more frequently changing between sister taxa in association with a phenotype.

We provided further evidence that this trend is true in general by calculating the phylogenetic signal of the top hits from each model using Ives and Garland’s *α* [37]. This statistic captures the rate of transitions between having and not having a binary trait (here, a gene) across a tree; higher values therefore correspond to more disagreement between closely related species and lower values correspond to more agreement. Indeed, across all four phyla, the linear model identified gene families with significantly lower Ives-Garland *α* than the phylogenetic model (Fig 2E, linear model *p <* 10^−16^), indicating that these genes’ presence versus absence tended to be driven more by clade-to-clade differences (i.e., shared evolution).

These results suggest that standard linear models can identify genes that are truly important for colonizing an environment, such as the healthy human gut, but in addition will identify other genes that may simply be common in clades associated with that environment. The latter set will likely include many false positive associations from the perspective of understanding functions necessary for living in the environment. Phylogenetic linear models overcome this problem by adding the expectation that closely-related species will have similar phenotypes and distantly-related species will have less similar phenotypes, effectively upweighting instances where this is not the case. These conclusions are supported by our simulations and by an *in vivo* functional screen (see section 2.6).

### 2.4 Gene families associated with gut prevalence provide insight into colonization biology

Several of the gene families that we observe to be associated with gut prevalence have previously been linked to gut colonization efficiency. For example, in Firmicutes, we noticed that several top hits were annotated as sporulation proteins (e.g., “Stage 0 sporulation two−component response regulator”, Fig 1C). Sporulation is known to be a strategy for surviving harsh environments (such as acid, alcohol, and oxygen exposure) that is used by many, but not all, members of Firmicutes. Resistance to oxygen (aerotolerance) is particularly important because many gut Firmicutes are strict anaerobes [38], sporulation is known to be an important mechanism of transmission and survival in the environment (reviewed in Swick et al. [39]), and sporulation ability has been linked to transmission patterns of gut microbes [31]. Our result associating sporulation proteins to gut prevalence provides further evidence for sporulation as a strategy that is generally important for the propagation and fitness of gut microbes.

In Bacteroidetes, we observed an association between gut prevalence and the presence of a pair of gene families putatively assigned to the GAD operon, namely, the glutamate decarboxylase *gadB* and the glutamate/gamma-aminobutyric acid (GABA) antiporter *gadC*. These genes show a complex pattern of presence that is strongly correlated with gut prevalence (Fig 2D, Fig S5). Results from research in Proteobacteria, where these genes were first described, shows that their products participate in acid tolerance. L-glutamate must be protonated in order to be decarboxylated to GABA; export of GABA coupled to import of fresh L-glutamate therefore allows the net export of protons, raising intracellular pH [40]. It was previously hypothesized that this acid tolerance mechanism allowed bacteria to survive the harshly acidic conditions in the stomach: indeed, if disrupted in the pathogen *Edwardsiella tarda*, gut colonization in a fish model is impaired [41]. *Listeria monocytogenes* with disrupted Gad systems also become sensitive to porcine gastric fluid [42]. However, while it has previously been shown that gut *Bacteroides* do contain homologs for at least one of these genes [40], their functional importance has not yet been demonstrated in this phylum. Our results provide preliminary evidence that this system may be important in Bacteroidetes as well as in Proteobacteria.

### 2.5 Using body sites as a control allows us to differentiate general dispersal from a specific gut advantage

The previous analyses have focused on modeling the phenotype of overall prevalence in the human gut. However, microbes could be prevalent in the gut for at least two main reasons. First, they could be specifically well-adapted to the human gut; second, they could simply be very common in the environment (i.e., highly dispersed). The presence or absence of a gene family could enhance either of these properties. Some genes might, for example, confer improved stress tolerance that was adaptive across a range of harsh conditions, while others might allow, for example, uptake and catabolism of metabolic substrates that were more common in the human gut than in other environments.

With this in mind, we analyzed the relative enrichment of microbes in the gut over other human body sites in 127 individuals from the Human Microbiome Project (HMP) study [33]. We chose other body sites as a control because the physical distance between sites within a host is much smaller than the distance between people, and microbes from one body site are likely to be commonly, if transiently, introduced to other body sites (e.g., skin to oral cavity). To find specifically gut-associated genes, we used the phylogenetic linear model to regress gene presence-absence on the logit-transformed conditional probability *P* (*e*_Gut_|*m*), i.e., the probability that a body site was the human gut given that a particular species *m* was observed, which we estimated using Laplace regularization (see Methods, 4.9). We identified 4,672 genes whose presence in bacterial genomes was associated with those species being present in the gut versus other body sites in at least one phylum (397 in Bacteroidetes, 1,572 in Firmicutes, 1,284 in Proteobacteria, and 1,507 in Actinobacteria).

Overall, the effect sizes for genes learned from this body site-specific model correlated only moderately with those learned from the “gut prevalence” models (median *R*^2^ = 0.06, range −0.06—0.24), indicating that these two quantitative phenotypes describe distinct phenomena. Additionally, the overlap between significant (*q* ≤ 0.05) hits for both models was small (median Jaccard index 0.054, range 0.011—0.089). These results are not surprising given that our regularized estimates of gut specificity were only moderately correlated with overall gut prevalence (Spearman’s *ρ* = 0.33, Fig S1), even when prevalence was calculated only from HMP gut samples (Spearman’s *ρ* = −0.15). This may arise from different genes being involved in dispersal or adaptation to many different environments versus those involved in adaptation specifically to the gut.

Indeed, when we compare enrichments for genes significant in either the body site or overall prevalence models alone (i.e., genes with *q* ≤ 0.05 in one model but *q >* 0.5 and/or wrong sign of effect size in the other), we observe large functional shifts (Fig 3). For example, in the gut prevalence model, but not the body site-specific model, Firmicutes were strongly enriched for “dormancy and sporulation” (*q* = 8.7 × 10^−7^). Because sporulation is likely useful in a wide range of environments beyond the gut, this result seems intuitive. Body site-specific results for Firmicutes were instead enriched for genes involved in “phosphate metabolism” (*q* = 0.12) and in particular the term “high affinity phosphate transporter and control of PHO regulon” (*q* = 0.05).

**Figure 3.**
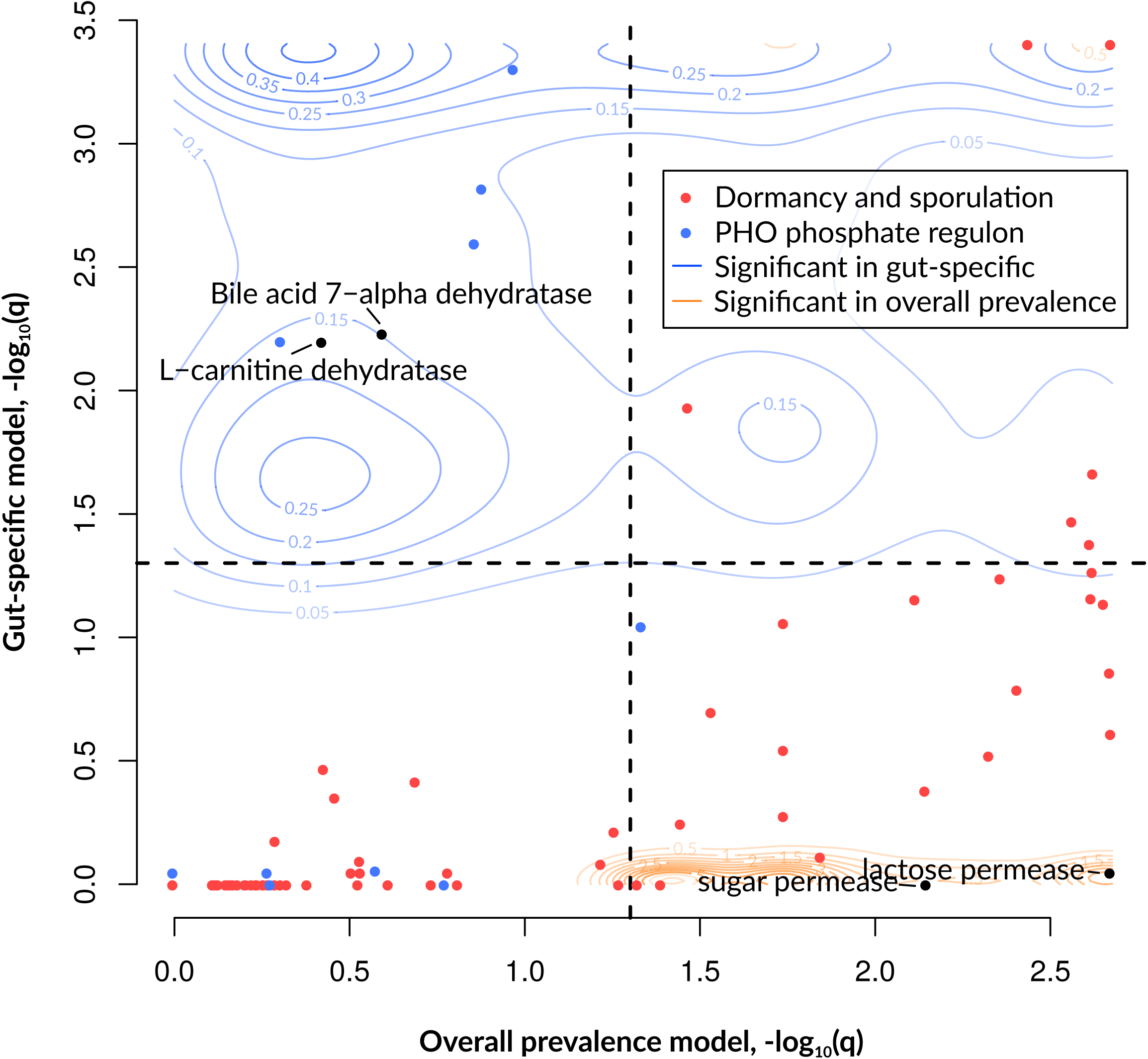
Comparison of results from the overall prevalence and body-site specific models for Firmicutes. FDR-corrected significance (as − log_10_(*q*)) of the overall model is plotted on the horizontal axis, whereas the same quantity for the body-site-specific model is plotted on the vertical axis. All FIGfams significant (*q* ≤ 0.05) in at least one of the two models are plotted as contour lines: FIGfams significant in the overall prevalence model (and possibly also the gut specific model) are plotted in orange, while FIGfams significant in the gut specific model (and possibly also the overall prevalence model) are plotted in blue. Selected SEED subsystems are displayed as colored points (legend), and selected individual genes are plotted as black points.

**Figure 4.**
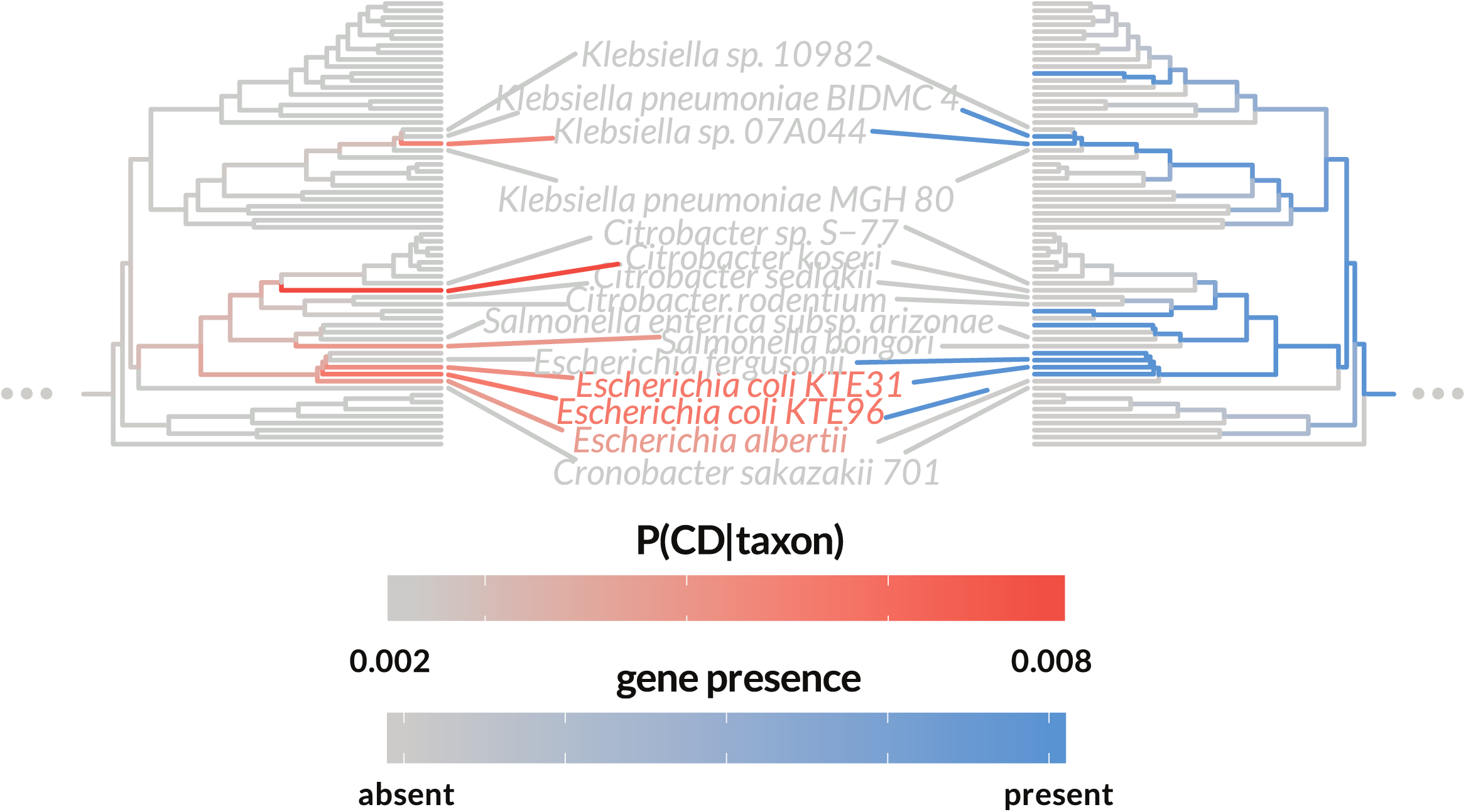
Genes involved in conjugative transfer are associated with Crohn’s disease-enriched species. The conjugation transcriptional regulator *traR* is plotted as an example. The left-hand tree is colored by each species’ disease specificity score, i.e., the conditional probability of Crohn’s given the observation of a given species (grey, which represents the prior, to orange, which represents a higher conditional probability). The right-hand tree is colored by gene presence-absence (grey, meaning absent, or blue, meaning present). The mirrored patterns drive the phylogeny-corrected correlation.

We also observed biologically-justified individual gene families that were significant in the body site-specific model but not the overall gut prevalence model. In Firmicutes, for example, carnitine dehydratase and bile acid 7-alpha dehydratase were both significant only in the body site-specific model, suggesting a specific role for these genes within the gut environment. Indeed, bile acids are metabolites of cholesterol that are produced by vertebrates and thus unlikely to be encountered outside of the host. While the metabolite L-carnitine is made and used in organisms spanning the tree of life, it is particularly concentrated in animal tissue and especially red meat, and cannot be further catabolized by humans [43], making it available to intestinal microbes. Bile acid transformation by gut commensals is a well-established function of the gut microbiome, with complex influences on health (reviewed in Staley et al. [44]).

In Bacteroidetes, we found that a homolog of the autoinducer 2 aldolase *lsrF* was significant only in the body site-specific model. Autoinducer 2 is a small signaling molecule produced by a wide range of bacteria that is involved in interspecies quorum sensing. The protein *lsrF*, specifically, is part of an operon whose function in *Escherichia coli* is to “quench” or destroy the AI-2 signal [45]. Further, an increase of the AI-2 signal has been shown to decrease the Bacteroidetes/Firmicutes ratio *in vivo* in the intestines of streptomycin-treated mice [46]. Degrading this molecule is therefore a plausible gut-specific colonization strategy for gut Bacteroidetes. These discovered associations make the genes involved, including many genes without known functions or roles in gut biology, excellent candidates for understanding how bacteria adapt to the gut environment.

### 2.6 Deletion of gut-specific genes lowers fitness in the mouse microbiome

Beyond finding evidence for the plausibility of individual genes based on the literature, we were interested in whether more high-throughput experimental evidence supported the associations we found between gut colonization and gene presence. To interrogate this, we used results from an *in vivo* transposon-insertion screen of four strains of *Bacteroides*. This screen identified many genes whose disruption caused a competitive disadvantage in gnotobiotic mice, as revealed by time-course high-throughput sequencing; 79 gene families significantly affected microbial fitness across all four strains tested [47]. Determining agreement with this screen is somewhat complicated by the fact that we associated gene presence to gut specificity across all members of the phylum Bacteroidetes, and not only within the *Bacteroides* genus. Significance of overlap therefore depends on what we take as the null “background” set, the cutoff used for significance, and the set of results from the screen we choose as true positives (Table S3).

Despite these complications, this analysis clearly showed that the 79 genes whose disruption led to lower fitness in the murine gut across all four *Bacteroides* species were over-represented among our predictions for gut-specific genes (odds ratio = 4.39, *q* = 8.3 × 10^−3^), and remained so if we only considered the gene families that were present in all *Bacteroides* species (odds ratio= 7.02, *q* = 3.0 × 10^−3^) (Table 1). Interestingly, we observed the opposite pattern for the overall prevalence model: the prevalence-associated genes we identified were actually depleted for genes found to be important *in vivo* (odds ratio = 0.18, *q* = 7.7 × 10^−3^). We believe that this is because the body-site-specific model, like the experiment, focused specifically on colonization efficiency, while the overall gut prevalence model would have included genes involved in persistence and dispersal in the environment and transfer between hosts. This experimental evidence supports the idea that environment-specific phylogenetic linear models truly identify genes that are important for bacteria to colonize an environment.

**Table 1.**
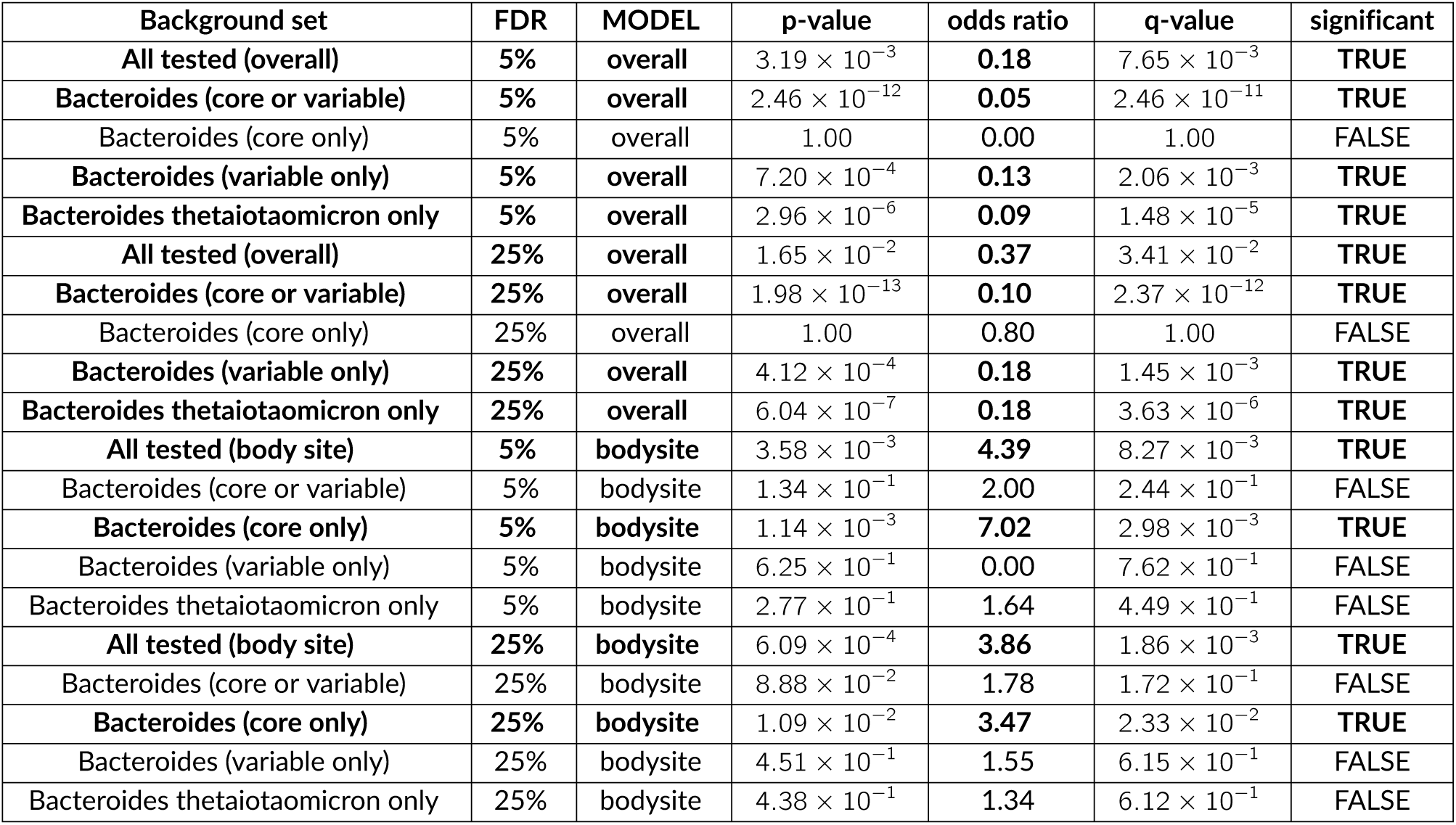
Assessment of agreement between the *in vivo* results from Wu et al. [47] and gut-specific (“bodysite”) vs. gut prevalence (“overall”) phylogenetic models. The background sets for enrichment tests were defined as follows: “all tested” (all gene families for which a phylogenetic model was fit), “Bacteroides (core or variable)” (all gene families with at least one representative in *Bacteroides* genome cluster pangenomes), “Bacteroides (core only)” (gene families that were present in all *Bacteroides* genome cluster pangenomes), “Bacteroides (variable only)” (gene families present in some but not all *Bacteroides* genomes clusters), and “Bacteroides thetaiotaomicron only” (only gene families present in *Bacteroides thetaiotaomicron*). The *p*-values are from Fisher’s exact tests. These comparisons have been excerpted from the full set, which can be seen in Additional Table S3; *q-*values were calculated based on this full set of tests using the Benjamini-Hochberg method [81].

### 2.7 We identify Proteobacterial gene families associated with microbes that are more prevalent in Crohn’s disease

The above analyses were performed with respect to the gut of healthy individuals from the mainly post-industrial populations of North America, Europe and China. However, we also know that taxonomic shifts are common between healthy guts versus the guts of individuals from the same population with diseases such as type 2 diabetes, colorectal cancer, rheumatoid arthritis, and inflammatory bowel disease (reviewed in Wang et al. [48]). One explanation for these results is that sick hosts select for specific microbial taxa, as with the links previously observed between Proteobacteria and the inflammation that accompanies many disease states [49]. Since gut microbes have also been implicated in altering disease progression (reviewed in Lynch and Pedersen [50]), identifying genes associated with colonizing diseased individuals may afford us new opportunities for intervention.

To identify microbiome functions that could be involved in disease-specific adaptation to the gut, we looked for genes that were present more often in microbes that discriminated case from control subjects. Specifically, we compared *n* = 38 healthy controls from the MetaHIT consortium to *n* = 13 individuals with Crohn’s disease [51, 52]. Similar to our analysis of gut versus other body sites, we used the conditional probability that a subject had Crohn’s disease *given* that we observed a particular microbe in their gut microbiome *P* (*e*_*CD*_|*M*) (see Methods). We identified 1,904 genes whose presence in bacterial genomes is associated with Crohn’s after correcting for phylogenetic relationships in at least one phylum (800 in Bacteroidetes, 272 in Firmicutes, 529 in Proteobacteria, and 319 in Actinobacteria).

Three of our top Proteobacterial associations were annotated as fimbrial proteins, including one predicted to be involved specifically in the regulation of type 1 fimbriae, or pili (FimE, association *q* = 4.0 × 10^−6^), cell surface structures involved in attachment and invasion. Crohn’s pathology has been linked to an immune response to invasive bacteria, and adherent-invasive *E. coli* (AIEC) appear to be overrepresented in ileal Crohn’s [53]. In an AIEC *E. coli* strain isolated from the ileum of a Crohn’s patient, type 1 pili were required for this adherent-invasive phenotype [54]. Chronic infection by AIEC strains was also observed to lead to chronic inflammation, and to an increase in Th17 cells and a decrease in CD8^+^ T cells similar to that observed in Crohn’s patients [55].

An additional striking feature of the results was the number of Proteobacterial proteins associated with greater risk of Crohn’s that were annotated as being involved in the the type III, IV, VI, and ESAT secretion systems (Fisher’s test *q* = 0.13). On further investigation, we found that these proteins were actually all predicted to be involved in conjugative transfer, a process by which gram-negative bacteria in direct physical contact share genetic material. More specifically, many of these genes were homologs of those involved in an “F-type” conjugal system for transferring IncF plasmids, which can be classified as a variety of type IV secretion system [56]. Previously, in a mouse model, gut inflammation was shown to stimulate efficient horizontal gene transfer in Proteobacteria by promoting blooms of *Enterobacteriaceae* and thus facilitating cell-to-cell contact [57]. Future work will be required to determine whether this increased conjugation is a neutral consequence of inflammation, a causative factor, or provides a selective advantage in the inflamed gut.

## 3 Discussion

The present analyses represent a first look into what can be learned by combining shotgun metagenomics with phylogenetically-aware models. Several extensions to our work could be made in the future. First, in addition to modeling prevalence, for instance, we could model abundance using a phylogenetic linear model with random effects [58], potentially allowing us to learn what controls the steady-state abundance of species in the gut. Additionally, we could also use these models to screen for epistatic interactions, which would be near-intractable even in systems with well-characterized genetic tools, but for which a subset of hypotheses could be validated by, e.g., comparing the fitness of wild-type microbes with double knockouts. While controlling the total number of tests would still be important to preserve power, an automated, computational approach to detecting gene interactions would still offer important savings in time and expense over developing a genome-wide experimental library of multiple knockouts per organism under investigation.

Currently, these analyses estimate species abundance and gene presence-absence from available sequenced isolate genomes. However, it has been estimated that on average 51% of genomes in the gut are from novel species [31]. Especially for case/control comparisons, using information from metagenomic assemblies could enable quantification of species with no sequenced representatives, and would yield a more accurate estimate of the complement of genes in the pangenome for species that do have sequenced representatives. This would be particularly helpful in gut communities from individuals in non-industrialized societies that are enriched for novel microbial species [31]. In fact, genes then could be treated as quantitative variables (e.g., coverage or prevalence) rather than binary, which is possible for covariates in phylogenetic linear models and simply changes the interpretation of the association coefficient *β*_1,*g*_.

Another potential extension would be to model prevalence and environment-specific prevalence for taxa other than the species clusters analyzed in this study. We focused on four prevalent and abundant phyla of bacteria, but our methods could be applied more broadly as long as quantitative phenotypes and genotypes could be accurately estimated. Phylogenetic linear modeling could also be applied directly to genera or higher taxonomic groups, although both phenotypes and genotypes would be averages over more diverse sets of genomes, which could result in associations with different signs canceling out. As more genome and metagenome data is generated for microbial populations over time, extensions of phylogenetic linear modeling (e.g., with random effects [58]) may also be useful for studying associations between phenotypes and evolving gene copy number and single nucleotide variants at the strain level. This application would require accurate trees with strains as leaves, each with estimates of a phenotype and genotype. Additionally, our current definition of species approximates a 95% average nucleotide identity (ANI) cutoff; while this approach is a standard bioinformatic approach [59], and appears to be a “natural boundary” in analyses of genome compendia [60], the precise definition of a bacterial species remains a matter of active debate, and in the future may include phenotypic information [61] or information about gene flow [62]. Beyond prevalence, other phenotypes will also be interesting to investigate, especially experimentally measured phenotypes from high throughput screens and other techniques that complement genomics.

In summary, using phylogenetic linear models, we were able to discover thousands of specific gene families associated with quantitative phenotypes calculated directly from data: overall gut prevalence, a specificity score for the gut over other body sites, and a specificity score for the gut in Crohn’s disease versus health. Importantly, we have shown through simulation and real examples that standard linear models are inadequate for this task because of an unacceptably high false-positive rate under realistic conditions. Furthermore, many of the results we found also have biological plausibility, both from the literature on specific microbial pathways and from a high-throughput *in vivo* screen directly measuring colonization efficiency. In addition to these expected discoveries, we also found thousands of novel candidates for understanding and potentially manipulating gut colonization. These results illustrate the potential of integrating phylogeny with shotgun metagenomic data to deepen our understanding of the factors determining which microbes come to constitute our gut microbiota in health and disease.

## 4 Methods

A graphical overview of our statistical methods can be found in Fig S2.

### 4.1 Species definition

We utilized the previously published clustering of 31,007 high-quality bacterial genomes into 5,952 species from the MIDAS 1.0 database [31] (http://lighthouse.ucsf.edu/MIDAS/midas_db_v1.0.tar.gz). These species clusters are sets of genomes with high pairwise sequence similarity across a panel of 30 universal, single-copy genes. The genomes in each species clustering have approximately 95% average genome-wide nucleotide identity, a common “gold-standard” definition of bacterial and archaeal species [63]. These species-level taxonomic units are similar to, but can differ from, operational taxonomic units (OTUs) defined solely on the basis of the 16S rRNA gene.

Taxonomic annotations for each species were drawn from the MIDAS 1.0 database. Some taxonomic annotations of species in the MIDAS database were incomplete; these were fixed by searching the NCBI Taxonomy database using their web API via the rentrez package [64] and retrieving the full set of taxonomic annotations.

### 4.2 Pangenomes

Pangenomes for all species used in this study were downloaded from the MIDAS 1.0 database. As previously described [31], pangenomes were constructed by clustering the DNA sequences of the genes found across all strains of each species at 95% sequence identity using UCLUST [65]. Pangenomes were functionally annotated based on the FIGfams [35] which were included in the MIDAS databases and originally obtained from the PATRIC [66] database. Thus, each pangenome represents the set of known, non-redundant genes from each bacterial species with at least one sequenced isolate.

### 4.3 Phylogenetic tree construction

The tree used for phylogenetic analyses was based on the tree from Nayfach et al. [31] based on an approximate maximum likelihood using FastTree 2 [67] on a concatenated alignment (using MUSCLE [68]) of thirty universal genes. Thus, each tip in the tree represents the phylogenetic placement for one bacterial species. For the current analyses, the tree was rooted using the cyanobacterium *Prochlorococcus marinus* as an outgroup, and the tree was then divided by phylum, retaining the four most prevalent phyla in the human gut (Bacteroidetes, Firmicutes, Actinobacteria, and Proteobacteria). One Actinobacterial species cluster, the radiation-resistant bacterium *Kineococcus radiotolerans* SRS30126, was dropped from the tree because it had an extremely long branch length, indicating an unusual degree of divergence. Finally, phylum-specific trees were made ultrametric using the chronos function in the R package ape [69], assuming the default “correlated rates” model of substitution rate variation. We performed this step because first, our taxa were contemporaneously sampled, and second, we assumed that our phenotypes of interest varied with divergence time, as opposed to the number of substitutions per site separating marker gene sequences [70].

### 4.4 Estimating species abundance across human associated metagenomes

Metagenome samples were drawn from subjects in the Human Microbiome Project (HMP) [33], the MetaHIT consortium [51, 52], a study of glucose control [71], and a study of type 2 diabetes [72]. Accession numbers were identified using the aid of SRAdb [73] and downloaded from the Sequence Read Archive (SRA) [74]. The relative abundance of bacterial species in the metagenomes was estimated using MIDAS v1.0 [31], which maps reads to a panel of 15 phylogenetic marker genes. Species relative abundances are computed as previously described [31] (“Species abundance estimation”): essentially, they are normalized counts of reads mapping to bacterial species, with non-uniquely mapped reads assigned probabilistically.

Accession IDs used can be found in Table S4. For prevalence estimates, we used healthy subjects from all four cohorts; for body site comparisons, we used only healthy subjects from HMP [33], and for the Crohn’s case-control comparison, we used only subjects from the MetaHIT consortium [51, 52].

### 4.5 Modeling gene-phenotype associations

The basic design is the same for all models that we fit: we model the effect of a categorical variable, gene (specifically, FIGfam family) presence vs. absence, on a particular phenotype estimated for many microbes from data.

Here, let 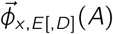 refer to a vector whose elements *ϕ*_*m,x,E*[,*D*]_(*A*) refer to an estimate of the phenotype *ϕ* for a microbe *m*, in an environment *e*_*x*_ from a set of *k* environments *E* = {*e*_1_, … *e*_*k*_}, optionally also adjusting for potential dataset effects *D*, based on a matrix of microbial presence-absence data *A*. We then model the effect on this phenotype of having vs. lacking each particular gene *g*, fitting one model per gene:

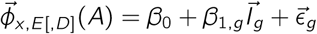

where *β*_0_ is a baseline intercept value, *β*_1,*g*_ is the effect size of gene *g*, 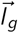 is a binary vector whose elements *I*_*g,m*_ are 1 when microbe *m*’s pangenome contains the gene *g* and 0 otherwise, and 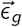 are the residuals. We then test the null hypothesis *H*_0_ : *β*_1,*g*_ = 0, yielding one *p*-value per gene; the resulting genewise *p*-values are finally corrected for multiple testing using an adaptive false discovery rate approach (*q*-value estimation).

The differences in the models we fit concern only how we obtain phenotype estimates 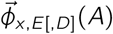, and our assumptions about how the residuals 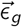 are distributed.

### 4.6 Fitting linear vs. phylogenetic models

The phylogenetic and standard linear models are very similar, except for the assumptions about the distribution of the residuals. In the standard linear model, the residuals are assumed to be independently and identically distributed as a normal distribution, i.e., *ϵ* _*g,m*_ ∼ *N*(0, *σ*^2^) or using multivariate notation 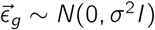. In the phylogenetic model, in contrast, the residuals are not independent: rather, they are correlated based on the phylogenetic relatedness of the species. They are therefore distributed *ϵ* ∼ *N*(0, σ), with the following covariance matrix:

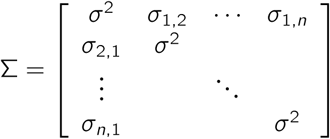

where *i* is the number of species, *σ*^2^ is the overall variance, and *σ*_1,2_ is the covariance between species 1 and species 2. Under the assumption of the phylogenetic model (evolution of a continuous phenotype according to Brownian motion), this covariance is proportional to the distance between the last common ancestor of species 1 and 2 and the root of the tree. Thus, very closely-related species have a common ancestor that is far from the root, while the last common ancestor of two unrelated species is the root node itself. This method was first described in Grafen [26]; for this study, we use the implementation in the phylolm R package [75].

*β*_1_ parameters were tested for a significant difference from 0 and the resulting *p*-values were converted to *q*-values using Storey and Tibshirani’s FDR correction procedure [76, 77].

### 4.7 Metagenomic presence-absence data

We use binary presence-absence data to calculate the phenotypes of interest. More formally, we conceptualize the metagenomic data as a matrix *A* of microbial presence-absence with dimensions *i* × *j*, where *i* is the number of microbes and *j* is the number of samples, and *a*_*m,n*_ is 1 if the relative abundance of microbe *m* (calculated using MIDAS’s taxonomic profiling [31]) is greater than 0, and 0 otherwise:

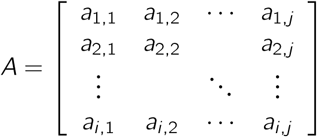

We conceptualize each *e*_1_, *e*_2_, …, *e*_*k*_ ∈ *E* as a set of indices, referring to samples collected from that environment, e.g., the oropharynx in healthy subjects, or the gut in subjects with Crohn’s disease, such that for all *e*_*x*_ ∈ *E, e*_*x*_ *⊆* {1, 2, …, *j*}. Because one environment may be tested in multiple studies, for our prevalence estimates, we also define a similar mapping of samples to datasets *d*_1_, *d*_2_, …, *d*_*l*_ ∈ *D* such that for all *d*_*y*_ ∈ *D, d*_*y*_ *⊆* {1, 2, …, *j*}. (For calculating environmental specificity scores, to avoid having to correct for unbalanced designs, we only use single datasets that measured all environments to be compared.) We also assume that *E* and *D* are partitions of {1, 2, …, *j*}, such that every sample is covered and no sample belongs to multiple *e*_*x*_ or *d*_*y*_

### 4.8 Estimating the prevalence phenotype

The first phenotype we consider is prevalence, *p*. Prevalence is usually defined as the fraction of samples in which a particular taxon is observed. Using the formulation above, the prevalence of microbe *m* in environment *e*_*x*_ and study *d*_*y*_ would be equal to:

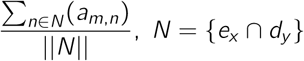

where we denote the quantity 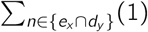, yielding the number of samples in environment *e*_*x*_, as ‖*N*‖.

We now take a slightly more general definition, such that a particular taxon’s true prevalence *p*_*m,N*_ is the probability of observing a particular microbe *m* in a set of samples *N, P* (*m*|*N*). Because *N* = *e*_*x*_ ∩ *d*_*y*_, we can also write this as *P* (*m*|*e*_*x*_, *d*_*y*_). More specifically, we can say that *p*_*m,N*_ is the probability parameter of the binomial random variable *a*_*m,N*_, which one can think of as generating the samples *a*_*m,n*_ such that *n* ∈ *N* in our matrix *A*:

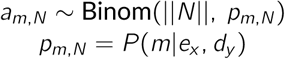

The maximum likelihood estimator of *p*_*m,N*_ given the data matrix *A* is then, as above, the fraction of subjects in *N* in which microbe *m* was observed:

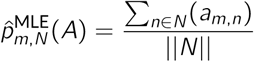

Because *p* is a proportion or probability, it is bounded between 0 and 1. The distribution of *p* is therefore highly non-normal, potentially violating assumptions of our regression model. We will therefore uselogit(*p*) as our phenotype. However, this now introduces a problem because 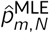 can take the values 0 and 1, leading to infinite estimates of logit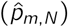. We therefore instead use a *shrunken* estimate of 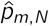.

Shrinkage estimators reduce the variance in the estimate of a parameter by combining it with prior information. These priors can be estimated from data (as in empirical Bayes approaches), estimated from independent information about the distribution of the parameter, or selected to be uninformative. Here, we use an uninformative prior, in this case a uniform distribution:

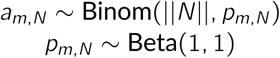

Mechanistically, this is equivalent to performing additive smoothing, which effectively adds one pseudocount to the numbers of absences and presences:

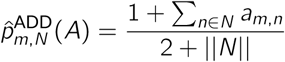

Finally, we note that this estimate of prevalence is only valid within a single study *d*_*y*_ ⊇ *N*. However, what we really want is an estimator of prevalence that depends only on the environment *e*_*x*_. We therefore marginalize out the effect of *d*_*y*_ :

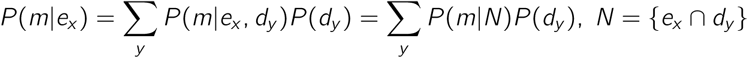

where we let the prior probability *P* (*d*_*y*_) simply be the probability of choosing a sample belonging to a dataset *d*_*y*_ out of all samples in environment *e*_*x*_, or:

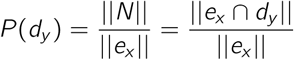

Effectively, this weights each dataset inverse-proportionally to the number of samples, so that the study with the largest number of samples does not dominate our estimates of prevalence. To avoid effects from additive smoothing dominating our estimates (as might happen if the same smoothing were applied to samples with different numbers of samples), we first obtain a marginalized version of the maximum-likelihood estimator, then perform additive smoothing on these marginalized estimates:

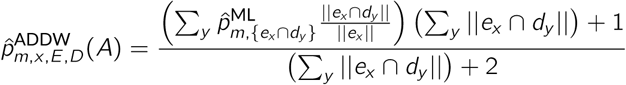

Finally, we use the logit of this estimate, i.e., 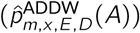, as the elements 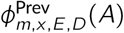 of our first phenotype 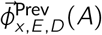:

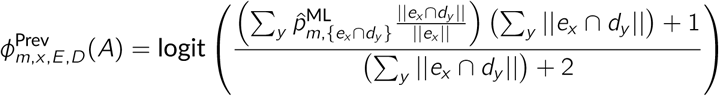

To recapitulate, we use a logit-transformed, shrunken estimate of prevalence in a given environment, weighted so that each study contributes equally.

### 4.9 Estimating environmental specificity scores

#### 4.9.1 Formulating the specificity score

Prevalence gives us information about how commonly a microbe is seen in a particular environment. While useful, this concept does not address the difference between microbes that are specific for a given environment and those that have a cosmopolitan distribution. We therefore wanted to design a statistic capturing this environmental specificity. We define this statistic in terms of how *predictive* a particular microbe is for one out of a set of possible environments. (For simplicity, we only consider cases in which all environments were measured within a single study, and therefore drop *D* from these equations. This estimator could be extended in the future to account for study effects as above.)

Recall that prevalence can be defined as the probability *P* (*m*|*e*_*x*_), i.e., the probability of observing microbe *m* in environment *e*_*x*_. To avoid potential sources of confounding error, we only consider environments both measured within the same study. We therefore let the environmental specificity score *s*_*m,x,E*_ equal the probability of observing a particular environment *e*_*x*_ out of a set of *k* environments *E* = *{e*_1_, *e*_2_, …, *e*_*k*_ *}*, given that we observe microbe *m*:

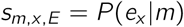

which, by application of Bayes’ rule and then marginalization, becomes:

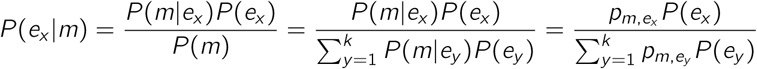

where *P* (*m*|*e*_*x*_) is the prevalence 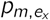 of microbe *m* in environment *e*_*x*_ ∈ *E*, and *P* (*e*_*x*_) is the prior probability of observing environment *e*_*x*_. The priors *P* (*e*) can be uninformative, in which case *P* (*e*_*x*_) = 1*/k* for all *x*, meaning that all environments are equally likely. This is the approach we take for body site comparisons. Alternatively, for a disease state, it could be drawn from actual epidemiological data about the frequency of that disease in the population of interest. This is the approach we take for the Crohn’s disease comparisons, taking *P* (*e*_CD_) = 0.002 [78], since in a Crohn’s case-control study, the fraction of individuals with Crohn’s will be much higher than the true prevalence of this disease in the population. In either case, the *P* (*e*_*x*_) values do not depend on the values in the dataset *A*. An intriguing third possibility that would depend on *A* would be to estimate the priors *P* (*e*_*x*_) based on the average observed *α*-diversity within environment *e*_*x*_, such that more diverse environments would be modeled as *a priori* more likely to contain any particular microbe.

#### 4.9.2 Motivating a shrunken estimator of *s*_*m,x,E*_ (*A*)

One simple way to estimate 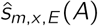 would be to simply plug in our estimates of 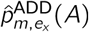, yielding:

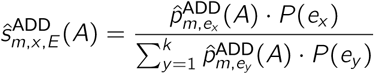

However, for cases in which the number of total observations of a microbe 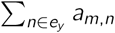 is low (imagine, e.g., a microbe that is observed once in environment *e*_1_ and zero times in *e*_2_), even the shrunken estimate 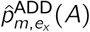 will have relatively high variance. This is particularly problematic here because both the numerator and denominator of 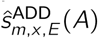 depend on 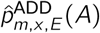, so as *p*_*m*,1,*E*_ → 0, the standard error of 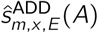 will tend to increase. This means that the microbes with the least-confidently estimated prevalences will tend to have high leverages in the regression, distorting the results. The confounding between the magnitude of 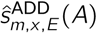 and its standard error also leads to heteroskedasticity, or unequal variance across the residuals, violating one of the main linear model assumptions.

To account for these issues, we construct a more aggressively-shrunk estimator of *s*_*m,x,E*_. We assume that most microbes do not differ substantially between environments, and therefore shrink estimates of *s*_*m,x,E*_ = *P* (*e*_*x*_|*m*) towards the prior probability *P* (*e*_*x*_), indicating that that this microbe is uninformative about the environment. To accomplish this, we use a maximum a posteriori (MAP) estimator, 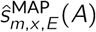, with a Laplace prior centered on *P* (*e*_*x*_). Laplace priors are also used in the Bayesian lasso to make parameter estimates more sparse, by shrinking them to zero. However, critically, we are not using the Laplace prior to perform model selection, since the exact same model is fit as in equation (1); we are only using it to reduce the variance in estimating *ŝ*_*m,x,E*_ ; unlike the Bayesian lasso, we therefore use no information about the independent variable (gene presence-absence) in obtaining estimates of *ŝ*_*m,x,E*_.

#### 4.9.3 An estimator of *s*_*m,x,E*_ using Laplace shrinkage

Before introducing this estimator, we briefly define the “environment-weighted prevalence” of a microbe *m* as its prevalence in each environment *e*_*y*_ *∈ E* weighted by the prior probability of that environment *P* (*e*_*y*_). Similarly to our previously-defined study-weighted prevalence estimator, this can be thought of as the overall probability of encountering a microbe, marginalized over environments:

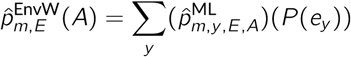

Since we are not taking the logit of this estimate, we can use the ML estimator. The environmental specificity score can then be modeled in this way:

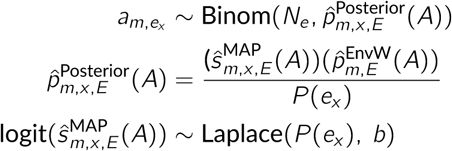

where *b* is a hyperparameter giving the width of the Laplace distribution, or equivalently the amount of shrinkage. Here, we are attempting to estimate *ŝ*^MAP^. The MAP estimate takes the following form:

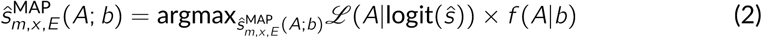

where *ŝ* is the parameter being estimated, *A* represents our data matrix, ℒ represents the likelihood function of the distribution from which the data is assumed to be drawn, *f* represents the density function of the prior distribution (without which the estimator reduces to the maximum-likelihood estimator), and *b* is the hyperparameter as above. We can expand this to give the final maximization:

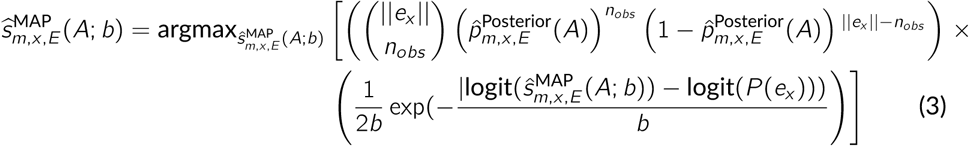

where we let 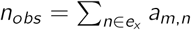, i.e., the number of times microbe *m* was observed in environment *e*_*x*_.

#### 4.9.4 Choosing the Laplace width parameter

To choose appropriate, dataset-specific values of *b*, which controls how much 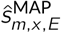(*A; b*) is shrunk back to the prior, we performed simulations. We chose a simulation-based approach instead of, for example, cross-validation because we lacked labeled examples of microbes that truly differed between environments. Instead, we constructed datasets *A*′ with elements 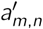 where we “knew” that some microbes (*m* ∈ *M*_0_) were not informative about the environment and others (*m* ∉ *M*_0_) had “true” differences, by simulating data with the following model:

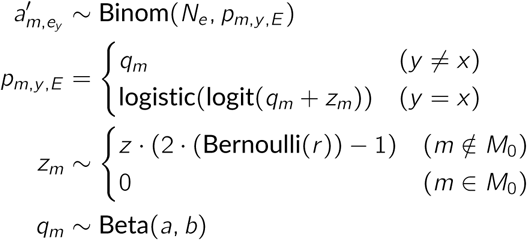

In other words, for each species *m* in different environments *e*_*y*_ *∈ E*, presence-absence 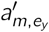 was modeled as a binomial random variable. The success parameter from this binomial was drawn from a Beta distribution with parameters *a* and *b*, which were fit from a single environment in the corresponding real dataset using maximum-likelihood, thus ensuring that the simulated species had similar baseline prevalences as real species. In species with no difference between environments *m* ∈ *M*_0_, the true prevalence *p*_*m,x,E*_ was set to be equal between the environment of interest *e*_*x*_ and all other environments; in species with true differences between environments (*m* ∉ *M*_0_), in contrast, the effect size *z* was either added or subtracted from the logit-prevalence (with the parameter *r* controlling the proportion of positive true effects). The number of null species ‖*M*_0_‖ was set to 25% of the total number of simulated species ‖*M*‖, which was matched to the real dataset.

For a given simulated dataset and value of *b*, the false positive rate (*F P R*_*b*_) and the true positive rates for *F >* 0 and *F <* 0 (*T P R*_*pos*_ and *T P R*_*neg*_, respectively) were calculated:

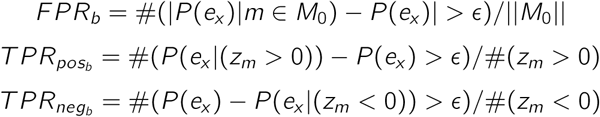

Since we are using numerical optimization, posterior probabilities are not always shrunk exactly to the prior; we therefore use a tolerance parameter *δ* set at *P* (*e*_*x*_) · 0.005 to account for numerical error. The tuning parameter *b* was then optimized according to the following piecewise continuous function, which increases from 0 to 1 until the false positive rate drops to 0.05 or lower (in order to guide the optimizer), and then increases above 1 in proportion to the average (geometric mean) of the positive and negative effect true positive rates:

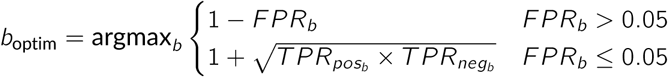

Given *z* = 2 and *r* = 0.5, for Crohn’s disease, *b*_optim_ was estimated at *b* = 0.14 and for the body site specificity, *b*_optim_ was estimated at *b* = 0.19. (Changing *z* to 1 or 0.5, or changing *r* to 0.1 or 0.9, resulted in very similar estimates of *b*_optim_. Additionally, *b*_optim_ estimates were consistent across several orders of magnitude of *E*; see Fig S4.)

#### 4.9.5 Worked example

An example showing the effect of this procedure on real data can be seen in Fig S3. The microbe *Bacillus subtilis* is detected once in the healthy cohort and once in the Crohn’s cohort, while *Bacteroides fragilis* is present in 24/38 healthy subjects but 13/13 Crohn’s subjects (Fig S3A). The maximum-likelihood values of 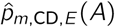 and *ŝ*_*m*,CD,*E*_ (*A*) are therefore much higher for *B. subtilis* than for *B. fragilis*, even though the evidence for a difference across environments in the prevalence of *B. subtilis* is much weaker (Fig S3B-C). In contrast, the Laplace prior (Fig S3D) successfully shrinks the estimate of the *B. subtilis* specificity score back to the baseline (*P* (*e*_CD_) = 0.002), while the evidence for *B. fragilis* overcomes this prior and yields an estimate close to the maximum-likelihood value (0.0031; Fig S3E).

### 4.10 Alternatives to shrinkage estimation of environmental specificity scores

An alternative to using the Laplace shrinkage estimator would be to allow all taxa to contribute to the regression, but to downweight taxa with less-confidently measured phenotypes. In generalized least squares (GLS), this is typically accomplished by scaling the variance-covariance matrix of the residuals by the variances of the estimators. This in effect says that the residuals are expected to be more dispersed around the regression line when the variance of the estimator is high: equivalently, this procedure weights each point in least-squares by the inverse of the estimator’s variance. We represent these variances as *v*_*m*_ ≡ (*SE*(*ŝ*_*m,x,E*_ (*A*)))^2^. The covariance matrix is then:

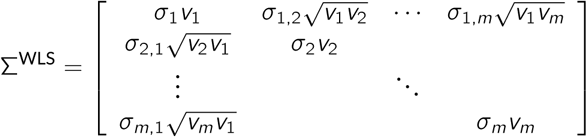

where *σ* values are as above. (Off-diagonal elements are weighted by the geometric mean of the variances, thus giving the same correlation structure as before.)

Because we have the data matrix *A* from which 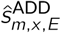 was estimated, we can estimate the variances of these estimates by bootstrapping. Denoting the *ŝ* estimates derived from bootstrap sample *c* as 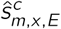, where *c ∈ {*1, *C}* and *C* is the number of bootstraps, and letting the mean across bootstrap samples be 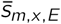, then 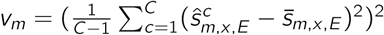.

Both approaches (Laplace shrinkage estimation and WLS) account for variability in the accuracy of estimating *s*_*m,x,E*_, but in different ways. We would expect them to agree more when *s*_*m,x,E*_ values were more confidently estimated (meaning the evidence for difference from the prior would be stronger and the extent of downweighting would be lower), and when more of the *s*_*m,x,E*_ values diverged substantially from the prior (leading to less sparsity in 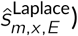). Indeed, when comparing body site specificities, which are based on higher numbers of samples and involve comparisons across highly divergent environments, both approaches yield more similar estimates, and become more concordant when the effect sizes are calculated only over significantly-different gene families. In contrast, for the Crohn’s disease comparison, in which the environments (healthy vs. diseased gut) are more closely related and the number of samples is smaller, the two methods tend to disagree more, especially in phyla where most taxa are shrunk back to the prior (Table S2). These results reflect the different underlying assumptions of each estimator: Laplace shrinkage assumes that taxa are not truly varying across environments without strong evidence, while WLS uses information from all taxa but downweights less-confidently-observed species. For the purposes of this manuscript, we focused on the results based on Laplace shrinkage estimation, since we believe the assumption that most taxa do not change is appropriate when comparing the same body site in health and disease. However, either approach may be preferable depending on the precise scenario being studied. It could also be possible to combine the two approaches by, for example, using the full posterior distribution of *ŝ*_*m,x,E*_ to derive weights.

### 4.11 Power analysis

To test the power and false positive rate of our method, we used parametric simulations, either under the null hypothesis in which a gene had no effect on the phenotype, or under the alternative hypothesis in which it had a defined effect. These involved generating one binary genotype and one continuous phenotype per simulation. These were parameterized as follows:

- The continuous phenotype 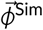 is simulated according to a Brownian motion model with parameters *β*_0_ corresponding to the ancestral state of the phenotype and *σ*^2^ corresponding to the phenotype’s overall variance (i.e., the diagonal of σ in the phylogenetic model).
- The binary genotype 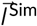 is generated from a Markov process as in Ives and Garland, with parameters *α* and *β*_1_. *α* gives the sum of the transition probabilities going from 0 to 1 and from 1 to 0. *β*_1_ gives the effect size: that is, how much the simulated phenotype influences the binary genotype (in logit space).

We perform the following process, given a choice of *α* and *β*_1_, for each phylum *h*:

1. Estimate the parameters 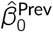 and 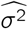 by fitting the following intercept-only phylogenetic model to the real prevalence phenotype:

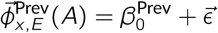

where *ϵ* ∼ *N*(0, σ) and Diag(σ) = *σ*^2^
2. For each of *B* simulations:
  a. Generate a continuous phenotype 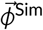 according to a Brownian motion process, evolving along the tree of phylum *h*, with parameters 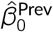 and 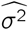.
  b. Generate a binary genotype 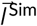 according to an Ives-Garland Markov process with parameters *α* and *β*_1_ and a covariate 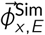.
  c. Fit the following regression equation:

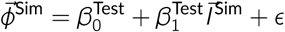

using either a standard linear model or the phylogenetic model (as specified in Methods 4.6).
  d. Return the *p*-value for the test of the null hypothesis 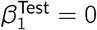.
3. The fraction of *p*-values ≤ 0.05 yields the power (when given *β*_1_ *>* 0) or the false positive rate (when given *β*_1_ = 0) of the test.

The binary genotype effect size *β*_1_ is not a linear function of the effect of the gene on prevalence. A more intuitive description of the effect size might be the (average) fold-change in prevalence associated with a gene’s presence. Because the parameters *β*^*Test*^ are on a logit scale, this quantity would be equal to:

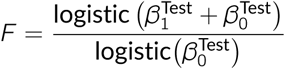

This quantity will depend on the amount of phylogenetic signal in the binary genotype *α*, the tree along which genotypes and phenotypes are simulated, and the “input” effect size *β*_1_. Accordingly, we simulated sets of 50 binary genotypes with given effect sizes *β*_1_ *∈* {0, 0.5, 0.75, 1.0, 1.25} and values of *α ∈* {0, 25, 50}. While there is substantial variation, in general an “input” effect size (i.e., *β*_1_) of 1.0 approximately corresponds to *F* ≈ 2, a two-fold difference in prevalence, and an “input” effect size of 0.5 corresponds to *F* ≈ 1.5, a 50% increase (Fig S8).

### 4.12 Assessing the potential impact of sampling with left-censoring

One potential pitfall with applying linear methods occurs when the distribution of the response variable (here, our phenotype) has a minimum value. This arises because, within a particular dataset, the lowest possible value of 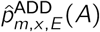 is equal to ‖*e*_*x*_‖^−1^, where ‖*e*_*x*_‖ is the number of samples in environment *e*_*x*_. This phenomenon is referred to as “left-censoring.” (Some may have encountered the term “left-censoring” in the context of participants who join a study having already experienced an event of interest. The time they experienced this event is therefore lower by an unknown amount than the lowest-possible measured value. While the domain and application are different, the statistical phenomenon is the same.) Left-censoring can result in inaccurate *p*-values because the variance is mis-estimated for the data points below the limit of detection. We therefore empirically assessed the impact of left-censoring in simulation, and also created extensions of the method to be used when its impact is noticeable.

Empirically, our prevalence phenotype 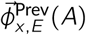 displays substantial left-censoring (Fig S6A). The distribution of this phenotype fits well to a normal with left-censoring at the limit of detection, in this case approximately −0.50 standard deviations below the mean (AIC using truncated normal and censoring: 20844.82; AIC using standard normal: 21833.38).

We therefore repeated the simulation process above, but after using our continuous phenotype to generate the binary genotype in step 2.b, we truncated the continuous phenotype 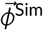 artificially at a specified number of standard deviations *K* below the mean, yielding 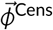. We then replaced 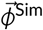 with 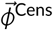 in the regression in step 2.c.

Using this simulation framework, we benchmarked three different ways to test the significance of 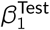 in the phylogenetic model. We computed p-values in one of the following three ways:

1. t-statistic: Return the *p*-value of a *t*-test of the null hypothesis that 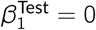, as in Methods 4.5.
2. Parametric Bootstrap: Simulate a number *C* of null binary genotypes 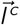 with *β*_1_ = 0, where *c ∈ {*1, …, *C*′*}*, and collect the estimates 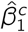 based on the following phylogenetic linear model:

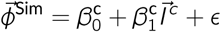

Then compute the fraction that are at least as extreme as the test statistic 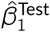 and return this as a *p-*value:

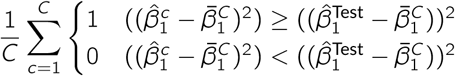

where 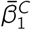 is the mean of 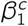 values. To save computation time on high *p*-values, we use early stopping: after every 25 such simulations, if the resulting *p*-value would already be guaranteed to exceed 0.05, we stop and return the *p*-value based on the current number of simulations.
3. Mock-Uncensored Bootstrap: As #2, simulate a number *C* of additional binary genotypes with *β*_1_ = 0. Instead of calculating *p*-values as in #2, however:
  a. Simulate the same number *C* of “uncensored” versions of the continuous phenotype 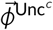, in effect “filling in” or imputing the censored values with random values from the predicted tail of the distribution (see Fig S6 B-C for an illustration):
    i. First, fit a left-truncated normal distribution to the part of 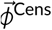 that is above the lowest value (assumed to be the limit of detection) by maximum likelihood (using `fitdistcens` in R package `fitdistrplus`), yielding mean *µ*^Trunc^, standard deviation 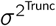, and lower truncation point 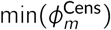 statistics (e.g., Fig S6B).
    ii. Next, for *c ∈ {*1, …, *C}*, generate a vector 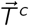 whose elements 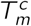 are realizations of the random variable 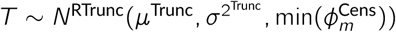. Here, *N*^RTrunc^ represents a right-truncated normal distribution, having mean *µ*^Trunc^, variance 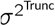, and upper truncation point 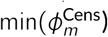. Then generate the “uncensored” vector 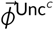 with elements 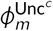 as follows:

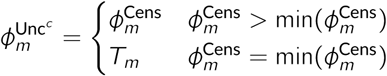

(An example can be seen in Fig S6C).
  b. For each *c ∈ {*1, …, *C}*:
    i. estimate a test statistic 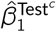 by fitting the following phylogenetic linear model:

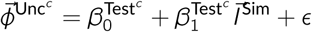
    ii. estimate a null test statistic 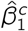 by fitting the following phylogenetic linear model:

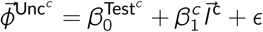
  c. Calculate *p*-values as follows:

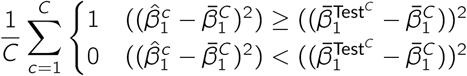

where 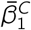 is the mean of the 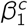 values and 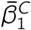 the mean of the 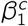 values.

Intuitively, the second method generates a null distribution via simulation, while the third method additionally reduces the impact of data points at the limit of detection, by randomly imputing them from the best-fit normal distribution. We then calculated power and FPR for each of these three methods, varying the amount of censoring *K* and the effect size *β*_1_ (see Fig S7). Interestingly, for the level of censoring in our data (*K* = −0.50), the false-positive rate in all three methods remained well-controlled, although power dropped. The mock-uncensored bootstrap had lower power overall and became more conservative, especially in the case where the phylogenetic signal was highest (*α* = 0) and where the level of censorship *K* was highest.

Another common approach to this problem is the tobit model: the true value of the response variable is treated as a hidden variable, and expectation-maximization is used to fit the regression parameters based on the observed censored values. Given that the degree of censoring we observed did not appear to inflate the false positive rate in any of the methods we tested, we opted not to construct a phylogenetic tobit model; however, this could be an interesting area of future research.

#### 4.1.3 Assessing the impact of compositionality

Because relative abundances are compositional (i.e., sum to 1.00), changes in highly abundant taxa combined with read sampling can lead to skewed estimates of relative abundance. For example, if a very abundant microbe exhibits large changes in relative abundance across samples, other microbes will appear to become less abundant simply because they make up a smaller proportion of the total reads, regardless of whether their level actually changes. This necessitates the use of compositional data analysis methods such as fitting intrinsically compositional distributions to the data (e.g., multinomial) or transforming it such that it is no longer compositional (e.g., the clr-transform [79]). However, the impact of these factors on prevalence was *a priori* less clear, because while prevalences are based on presence-absence, which could be affected by sampling, prevalences themselves do not have to sum to 1.

Let *R* be an *i*′ × *j* matrix of read counts whose elements *r*_*m,n*_ correspond to the number of reads mapping to the single-copy marker genes for microbial species *m* in sample *n*, where *i*′ is the number of microbes with at least one read in one of the *j* samples.

We resampled *R* using Dirichlet-multinomial sampling as follows. We first determined the set of microbes *M*^*Y*^ that have at least one read across our *j* samples but are not among the *Y* -most abundant microbes (so *M*^100^ would exclude the 100 most-abundant species). We then constructed a Dirichlet distribution *u*_*n*_ from each 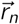 such that:

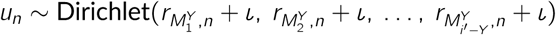

where *ι* is a pseudocount corresponding to the average number of reads for the average microbe:

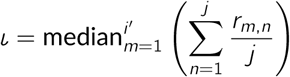

We then drew a multinomial distribution *v*_*n*_ from each *u*_*n*_, then based on this multinomial distribution, drew a count vector 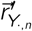 with total number of counts Σ*m r*_*m,n*_. These count vectors constituted the *columns* of a resampled dataset 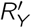, with elements obtained by Dirichlet-multinomial sampling from the original dataset *R*, after excluding the *Y* most prevalent microbes.

First, we examined how increasing *Y* impacted estimates of relative abundance, that is, how much relative abundance profiles were distorted by the effects of sampling in the presence of species with large read counts. We transformed the resampled “read count” dataset 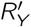 to a matrix of relative abundances 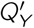 by dividing each column 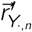 *by* 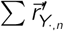 to yield normalized relative abundances 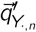. We then applied the clr-transform to correct for compositionality effects, yielding a matrix of transformed abundances 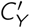 with columns:

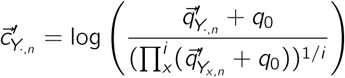

where *q*_0_ is the minimum non-zero value of 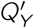, i.e., 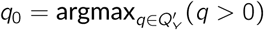.

After applying the clr-transform to compensate for compositionality artifacts, we tested the correlation of the row vectors 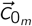. with the corresponding 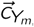. As expected, the median correlation was low (for Y=10, *r*_Median_=0.21) and continued to decrease as Y increased from 10 to 100 (*r*_Median_=0.16) (Fig S9).

In comparison, we next examined how increasing *Y* impacted prevalence. We first transformed each the resampled read matrix 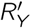 into a binary presence-absence matrix 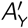, then calculated 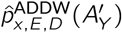 for each value of *Y*. Finally, we calculated the correlation of 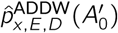 to each 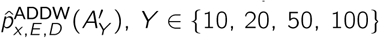, *Y ∈ {*10, 20, 50, 100*}*. Strikingly, we found that prevalences were highly correlated even out to *Y* = 100, where *r* = 0.91 (Fig S9).

These analyses suggest that unlike relative abundance profiles, our estimates of prevalence were robust to sampling and compositionality effects S9.

### 4.14 Enrichment analysis

Enrichment analysis was performed using SEED subsystem annotations for FIGfams [35, 80]. Each subsystem was tested for a significant overlap with significant hits from the linear models (*q* ≤ 0.05), given the set of FIGfams tested, by Fisher’s exact test. For each gene set, a 2 × 2 contingency table was constructed with the following form:

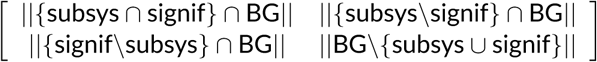

where “subsys” is the set of FIGfams in a given SEED subsystem, “signif” is the set of FIGfams in a particular phylum that were significant hits, and “BG” is the set of all FIGfams tested in that phylum. Two-tailed *p*-values were corrected using the Benjamini-Hochberg procedure [81] and an FDR of 25% was set for detecting significant enrichment and depletion (only enrichment is reported). We used this significance threshold in accordance with accepted practice for gene set enrichment analysis [82]. We used the Benjamini-Hochberg procedure since unlike the *q*-value method it does not require the estimation of the proportion of true nulls, which is more difficult with small numbers of tests.

### 4.15 Overlap with *in vivo* results

Results of the screen were obtained from the Supplemental Material of Wu et al. (downloaded on 2017 May 3) [47]. Genes were mapped to FIGfams by matching identifiers in the Supplemental Material to genome annotations from PATRIC [66]. Significance of overlap between these genes and the results for the Bacteroidetes phylum from the body-site-specific or overall models was determined by Fisher’s exact test.

This test depends both on how we determine which genes from the *in vivo* screen count as true positives, and on the choice of the “background set,” i.e., which genes would be possible to find in the *in vivo* study. Rather than committing to one method of picking the “true positive” and “background” sets, we instead enumerated several possibilities, performed all possible combinations (Table S3), and corrected for multiple comparisons. The options we tested for true positive sets were 1. genes in the screen that were significantly associated with fitness in all four *Bacteroides* strains tested, 2. *Bacteroides thetaiotaomicron* genes significantly associated with diet-independent fitness effects, and 3. *B. theta* genes associated with either diet-dependent or -independent effects. The background sets we tested were 1. all gene families for which a phylogenetic model was fit, 2. all gene families appearing at least once in a *Bacteroides* genome cluster pangenome, 3. all gene families present in all *Bacteroides* pangenomes, 4. gene families present in some but not all *Bacteroides* pangenomes, and 5. gene families present in *Bacteroides thetaiotaomicron.* Similarly to our approach to gene set enrichment analysis, for each test we assembled a 2×2 contingency table as follows:

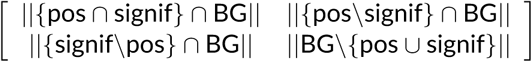

where “pos” refers to the true positive FIGfam set, “signif” refers to the set of significant FIGfam hits from the phylogenetic model, and “BG” refers to the background FIGfam set. The full results are depicted in Table SS3, with the results for true positive set #1 excerpted from this full comparison in Table 1.

### 4.16 Codebase

The code used to perform these analyses is available at http://www.bitbucket.com/pbradz/plr in the form of an Rmarkdown notebook.

### 4.17 Glossary of notation

- Data and metadata
  - *A*: *i* × *j* binary matrix of microbial presence-absence, where *i* is the number of microbes, *j* is the number of samples, and *a*_*m,n*_ is 1 when microbe *m* is observed in sample *n* and 0 otherwise
  - *a*_*m,N*_ : a vector of presence-absences for microbe *m* in samples *n ∈ N*
  - *e*_*x*_, *d*_*y*_ : environment *x* or study population *y*, each corresponding to a set of samples
  - *E* = {*e*_1_, …, *e*_*k*_}: the set of environments being studied or compared (e.g., body sites; health vs. disease)
  - *D* = *{d*_1_, *…, d*_*l*_ *}*: the set of study populations
- Linear models
  - 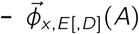: phenotype vector, calculated for environment *e*_*x*_ ∈ *E* and potentially adjusting for dataset effects *D*, based on microbial presence-absence matrix *A*, with elements corresponding to phenotype estimates for individual microbes
  - 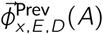: prevalence phenotype estimates (based on 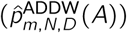; see below).
  - 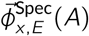: environmental specificity score phenotype estimates (based on logit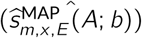; see below).
  - *β*_0,*g*_: in the linear model for gene *g*, intercept term used to model the average value of a given phenotype 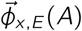
  - *β*_1,*g*_: in the linear model for gene *g*, the effect of having vs. not having gene *g* on a given phenotype 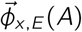
  - 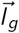: the binary vector of gene presence-absence whose elements are *I*_*g,m*_, equal to 0 if the gene *g* is absent in microbe *m* and 1 if it is present
- Phenotype estimation
  - *p*_*m,N*_ : prevalence, the probability of observing a microbe *m* in a set of samples *N P* (m|N)
  - 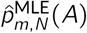: the maximum-likelihood estimate of prevalence, based on the presence-absence matrix *A*
  - 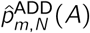: an estimate of prevalence based on the presence-absence matrix *A* using additive smoothing
  - 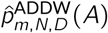: an estimate of prevalence based on the presence-absence matrix *A* using additive smoothing, and additionally weighting by the inverse number of samples per dataset in *D*
  - 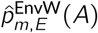: an estimate of the prevalence across environments, weighted by their probability (i.e., *P* (*m*) obtained by marginalizing *P* (m|e_x_))
  - *s*_*m,x,E*_ : environment specificity, the probability of being in a particular environment *e*_*x*_ given that microbe *m* was observed *P* (*e*_*x*_|*m*)
  - *ŝ*_*m,x,E*_ (*A*): an estimate of environment specificity based on presence-absence matrix *A*
  - *b*: a hyperparameter controlling the width of the Laplace prior on *ŝ*_*m,x,E*_ (*A*) (i.e., the amount of shrinkage in the estimate)
  - *b*_optim_: a value of *b* optimized for sensitivity and specificity in parametric simulations
  - *P* (*e*_*x*_): the prior probability of encountering environment *e*_*x*_ ; we use either an uninformative uniform prior (for bodysites), or take this prior from epidemiological data (for disease comparisons)
  - 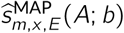: a maximum a posteriori (MAP) estimate of environment specificity score for environment *e*_*x*_ based on presence-absence matrix *A* and the shrinkage hyperparameter *b*; in this paper we calculate environmental specificity scores for *x* = CD (Crohn’s disease specificity) and *x* = Gut (healthy gut specificity)
- Simulation and censoring analysis
  - 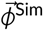: a simulated continuous phenotype
  - 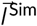: a simulated binary genotype (gene presence-absence)
  - *α*: Ives-Garland *α*, the sum of the transition probabilities from 0 to 1 and from 1 to 0 in a Markov model of binary trait evolution across a tree (i.e., a measure of phylogenetic signal in a binary trait)
  - *β*_0_: assigned parameter giving the ancestral state of the simulated genotype 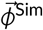
  - *β*_1_: assigned parameter giving the degree to which the continuous phenotype 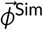affects the binary genotype 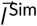; a measure of effect size of *phenotype on gene*
  - 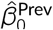: the estimated ancestral state of our prevalence phenotype 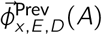, using a Brownian motion model of trait evolution
  - 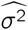: the estimated (non-phylogenetic) variance of our prevalence phenotype 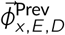(A)
  - 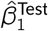: estimated value of effect of *gene on phenotype* from the phylogenetic linear model
  - *F* : ratio of prevalences, comparing taxa with a given gene (numerator) to taxa without (denominator); an alternative measure of effect size of *gene on phenotype*
  - 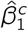: when using a bootstrap null, estimated value of effect of *null gene on phenotype* from the phylogenetic linear model
  - 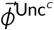: a version of the phenotype 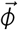 where values at the certain limit of detection have been imputed based on a truncated normal
  - 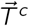: a vector the same length as 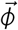 whose elements have been randomly drawn from a truncated normal distribution
  - 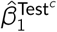: an estimated value of effect of *gene on the mock-uncensored phenotype* 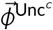
  - *K*: the value at which left-censoring starts for a phenotype 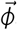, expressed as standard deviations below the mean
- Compositionality with resampling analysis
  - *R*: an *i*′ × *j* matrix with elements *r*_*m,n*_ corresponding to the number of reads mapping to microbe *m* in sample *n*
  - *M*^*Y*^ : a set of microbes with at least one read in *R*, excluding the top *X*-most abundant microbes
  - *ι*: pseudocount used in constructing Dirichlet distributions of microbial relative abundance
  - *u*_*n*_: a Dirichlet distribution fit to sample *n*
  - *v*_*n*_: a particular draw from a Dirichlet distribution representing a multinomial distribution, from which resampled read counts are drawn
  - 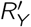: a matrix of resampled read counts with dimensions (*i*′− ‖*Y*‖) × *j*, where read counts are drawn from *v*_*n*_
  - 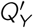: a matrix of resampled relative abundances derived from 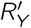 such that every column adds up to 1
  - 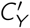: a matrix of clr-transformed relative abundances derived from 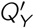
  - *q*_0_: a pseudocount used in the clr-transformation

## 5 Author contributions

- Patrick H. Bradley. Roles: Conceptualization, Data Curation, Formal Analysis, Investigation, Methodology, Software, Validation, Visualization, Writing – Original Draft Preparation, Writing – Review & Editing
- Stephen Nayfach. Roles: Data Curation, Software, Resources, Writing – Review & Editing
- Katherine S. Pollard. Roles: Conceptualization, Methodology, Funding Acquisition, Project Administration, Supervision, Writing – Original Draft Preparation, Writing – Review & Editing

## 7 Acknowledgements

The authors would like to thank Joshua Ladau, Nandita Garud, and other members of the Pollard and Turnbaugh groups, attendees of the 2017 Keystone meeting on the Microbiome in Health and Disease and attendees of the Second Workshop in Statistics and Algorithmic Challenges in Microbiome Data Analysis (SACMDA2), and the anonymous reviewers for helpful suggestions and discussions.

---

**Table S2.**
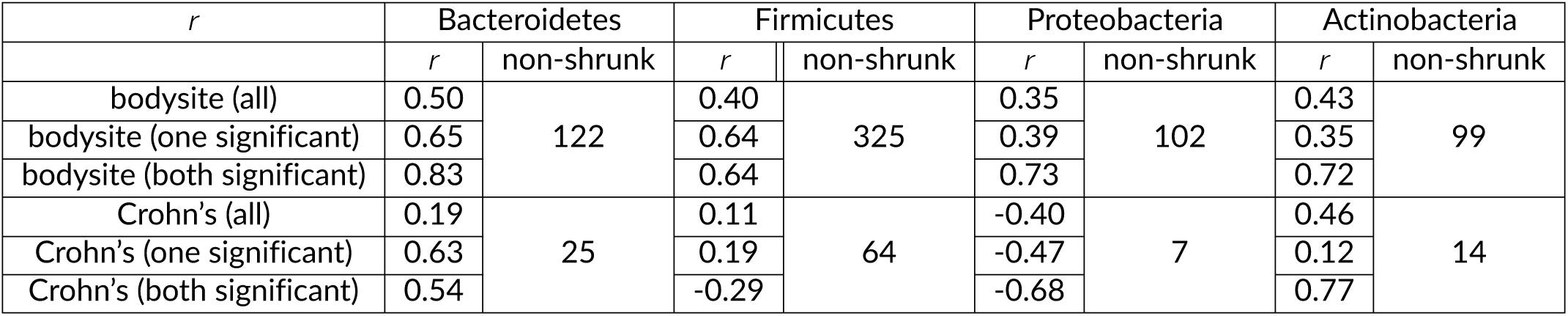
Concordance of *β*_1,*g*_ estimates for weighted least squares vs. Laplace shrinkage procedures. Pearson’s correlation coefficient *r* comparing estimates of *β*_1,*g*_ for the weighted phylogenetic least squares procedure with the unweighted phylogenetic least squares using Laplace-shrunken estimates of specificity scores were computed across: all tested genes (all), all genes significant in either the weighted or shrunken phylogenetic model (one significant), or all genes significant in both models (both significant). Comparisons were performed for both body site environmental specificity scores (bodysite) and Crohn’s disease (Crohn’s). Additionally, the number of taxa not shrunk back to the prior by Laplace shrinkage for each environmental specificity score are given (non-shrunk).

| $r$ | Bacteroidetes | | Firmicutes | | Proteobacteria | | Actinobacteria | |
| --- | --- | --- | --- | --- | --- | --- | --- | --- |
| | $r$ | non-shrunk | $r$ | non-shrunk | $r$ | non-shrunk | $r$ | non-shrunk |
| bodysite (all) | 0.50 | 122 | 0.40 | 325 | 0.35 | 102 | 0.43 | 99 |
| bodysite (one significant) | 0.65 |  | 0.64 |  | 0.39 |  | 0.35 |  |
| bodysite (both significant) | 0.83 |  | 0.64 |  | 0.73 |  | 0.72 |  |
| Crohn's (all) | 0.19 | 25 | 0.11 | 64 | -0.40 | 7 | 0.46 | 14 |
| Crohn's (one significant) | 0.63 |  | 0.19 |  | -0.47 |  | 0.12 |  |
| Crohn's (both significant) | 0.54 |  | -0.29 |  | -0.68 |  | 0.77 |  |

## Supporting Information

**Table S1. Species prevalences, gut specificities, and Crohn’s disease specificities for all genome clusters (species) tested.** logit.Prevalence, logit.BodySite, and logit.Crohns column titles refer to our estimates of 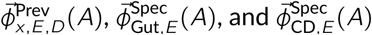, respectively. Row labels (5-digit numbers) correspond to MIDAS taxon IDs.

**Table S3. Full assessment of whether genes linked to microbial fitness in an *in vivo* experiment [47] were enriched for significant hits of the body site-specific and overall gut prevalence models.** The different sets of true positives were defined as: “Bacteroides” (genes in the screen significantly associated with fitness in all four strains), “BthetaDietIndep” (genes present in *Bacteroides thetaiotaomicron* that had diet-independent fitness effects in the screen), and “BthetaAny” (same, but for diet-dependent as well as -independent effects). The “background sets” were defined as follows: “all tested” (all gene families for which a phylogenetic model was fit), “Bacteroides (core or variable)” (all gene families with at least one representative in *Bacteroides* genome cluster pangenomes), “Bacteroides (core only)” (gene families that were present in all *Bacteroides* genome cluster pangenomes), “Bacteroides (variable only)” (gene families present in some but not all *Bacteroides* genomes clusters), and “Bacteroides thetaiotaomicron only” (only gene families present in *Bacteroides thetaiotaomicron*). Two false discovery rates for each model were tested (5% and 25%). Fisher tests yielded *p-*values that were then converted to *q-*values using the Benjamini-Hochberg approach [81].

**Table S4. SRA accession IDs used to estimate prevalence and environmental specificity scores.**

**Figure S1.**
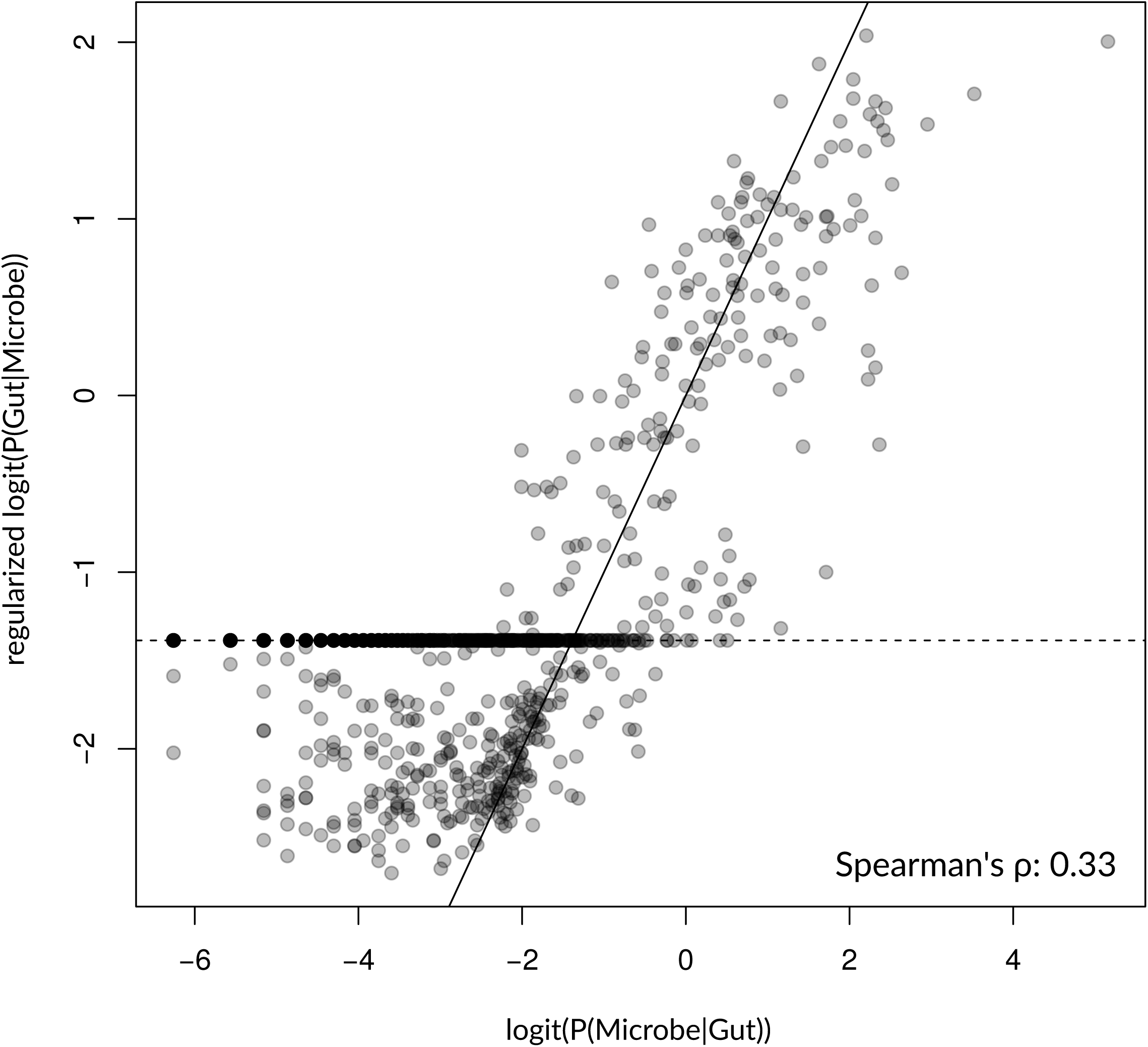
Estimates of logit-gut prevalence (x-axis) vs. logit-gut environmental specificity score (y-axis), showing only modest correlation.

**Figure S2.**
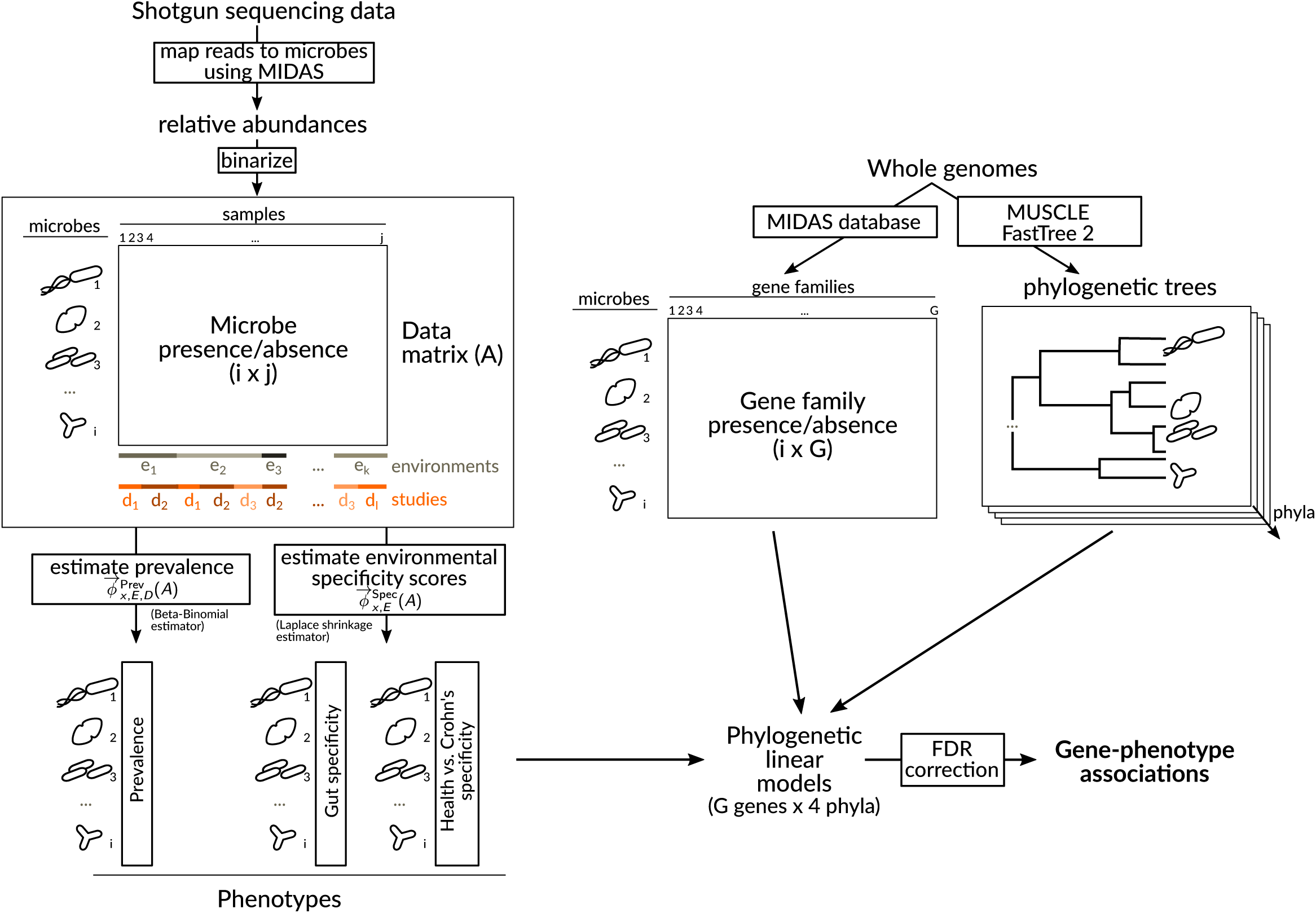
Method overview. Using MIDAS, we calculate species relative abundances from shotgun sequencing data. These are binarized to yield a matrix of microbial presence/absence, with rows corresponding to microbes and columns corresponding to samples. Samples are organized into environments (i.e., the environments from which the sample was collected) and into datasets (corresponding to samples collected as part of the same project). Using the presence/absence matrix together with these metadata, we estimate phenotype vectors 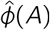, whose elements are estimates of microbial phenotypes. These phenotypes fall into two groups: prevalence 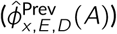 and environmental specificity scores 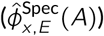. Separately, we use the whole genomes incorporated into the MIDAS database to assemble a matrix of gene presence/absence in the pangenome of microbes, and to construct a phylogenetic species tree based on previously-validated single-copy marker genes. We subset this tree to yield four phylum-specific trees. The inputs to our phylogenetic models are a phenotype vector, a gene presence-absence vector, and a phylogenetic tree. Based on these models, we estimate *p*-values for a non-zero effect of the gene on the phenotype, then convert these *p*-values into *q*-values to obtain predicted gene-phenotype interactions at a given false discovery rate (here, 5%).

**Figure S3.**
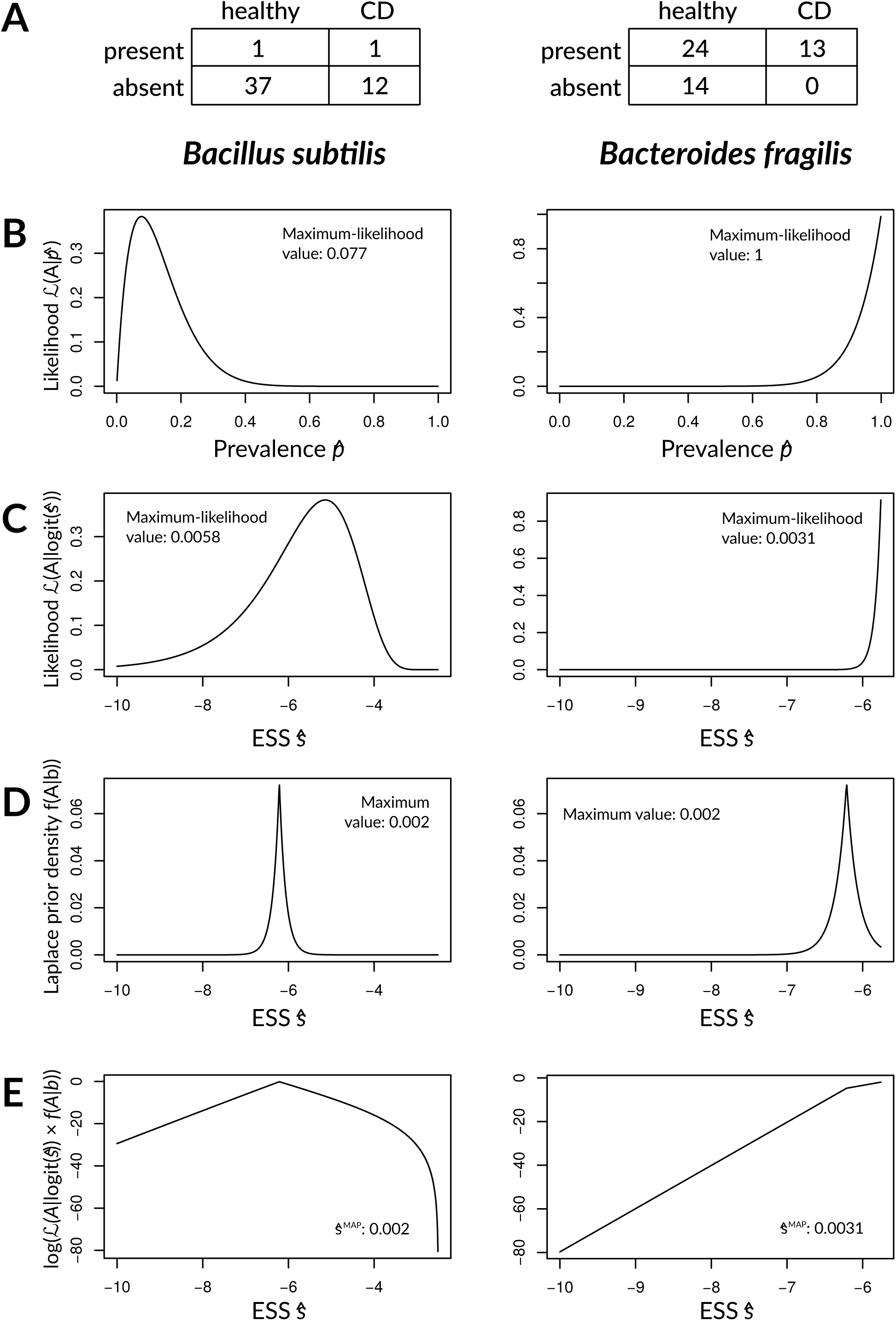
Laplacian regularization reduces noise in estimating *ŝ*_*m*,**CD**,*E*_ (*A*). Two species are compared, one that was infrequently observed in both Crohn’s disease cases and controls (*Bacillus subtilis*, left) and one with a significant bias for Crohn’s disease cases (*Bacteroides fragilis*, right). A) Total counts across subjects for *Bacillus subtilis* and *Bacteroides fragilis*. B) Likelihood function for *ŝ*_*m*,CD,*E*_ (*A*), or prevalence in Crohn’s disease. The maximum-likelihood value is given in the inset. C) Unregularized likelihood for logit(*ŝ*_*m*,CD,*E*_ (*A*)), or the environmental specificity of the microbe. Note that the maximum-likelihood value (inset) was actually almost twice as large for *Bacillus subtilis* as for *Bacteroides fragilis* despite the relative paucity of data for *B. subtilis* (compare Y-axes, which show that the distribution for *B. subtilis* is flatter). D) Laplace prior around *P* (*e*_CD_) = 0.002 with width parameter *b* = 0.15 (optimized using simulation). E) Log-likelihood plot for the posterior 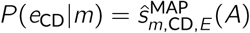, obtained by taking the product of the prior distribution and the unregularized distribution. The maximum *a posteriori* (MAP) estimates are the modes of these distributions (inset).

**Figure S4.**
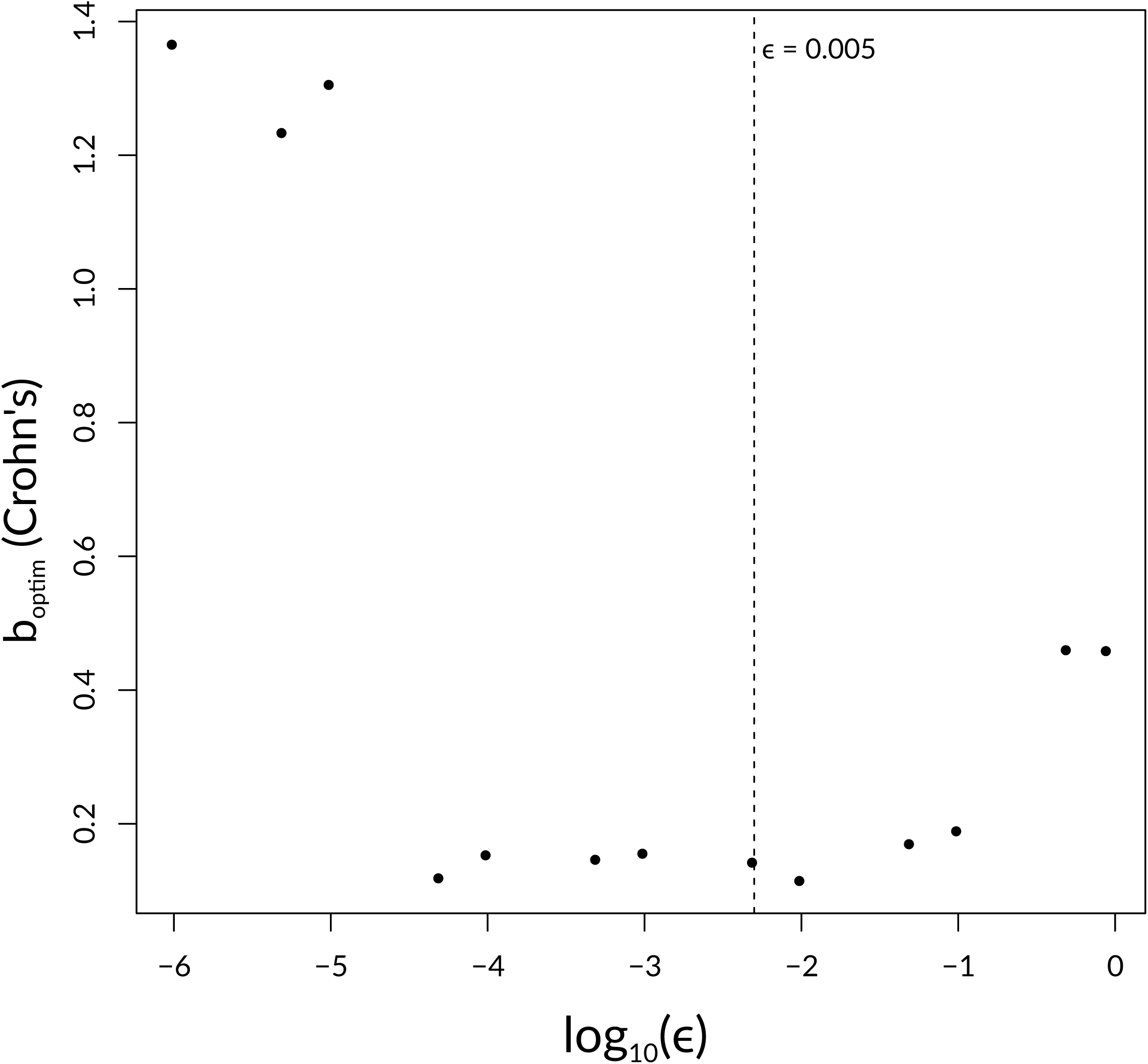
Sensitivity plot for *E* tolerance parameter in Laplace shrinkage. Y-axis gives the best *b*_optim_ value obtained given a particular log_10_(*ϵ*) selected when performing Laplace shrinkage of 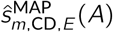 estimates. The value of *E* used in the manuscript (0.005) is highlighted with a vertical dashed line.

**Figure S5.**
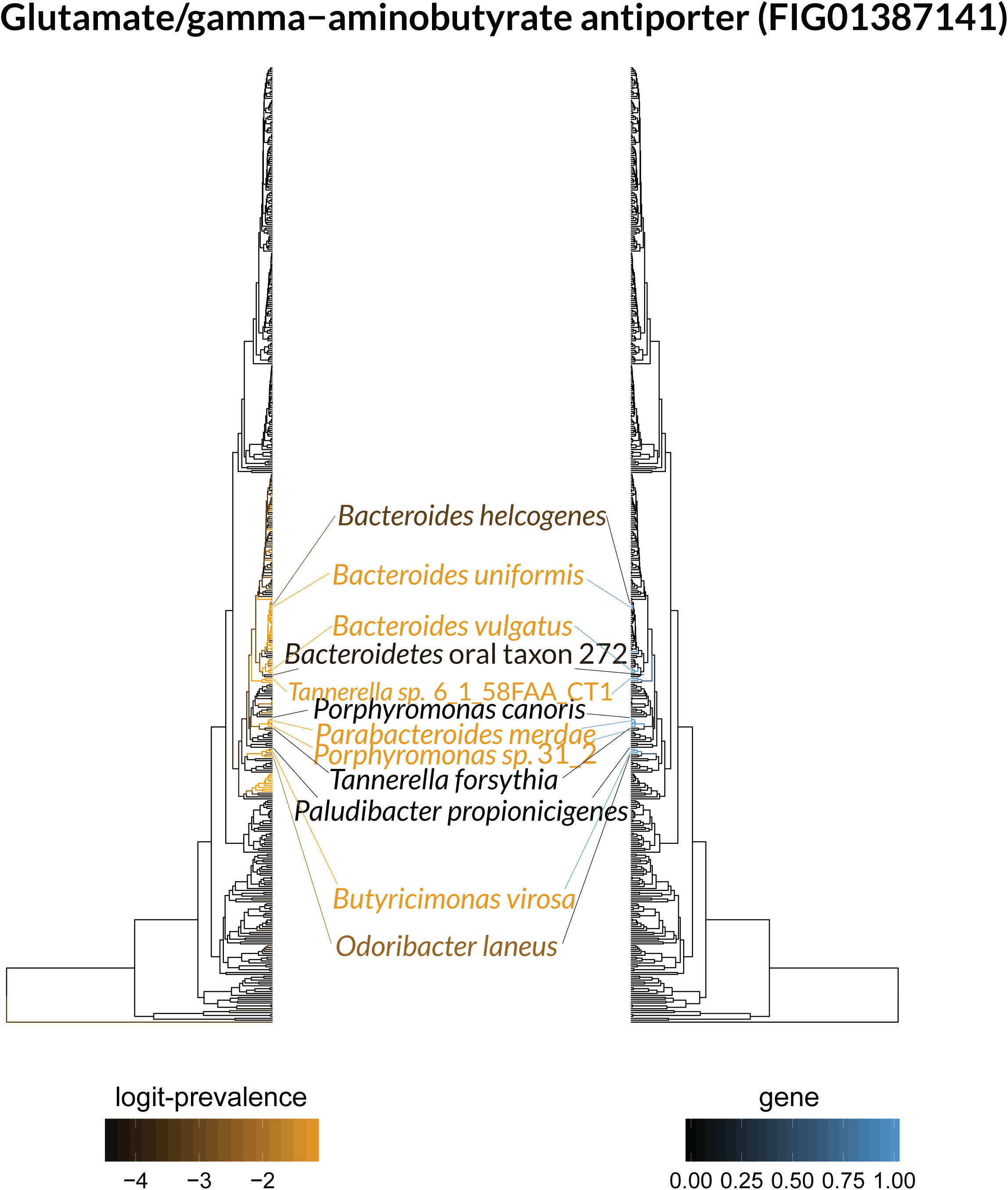
Illustration showing logit-prevalence vs. the pattern of glutamate-GABA decarboxylase (*gadB*) inheritance in Bacteroidetes. As in Fig 2, the tree on the left is colored by species prevalence (black to orange), while the tree on the right is colored by gene presence-absence (blue to black), with selected species called out in the middle, and lines linking species labels to leaves that match leaf color.

**Figure S6.**
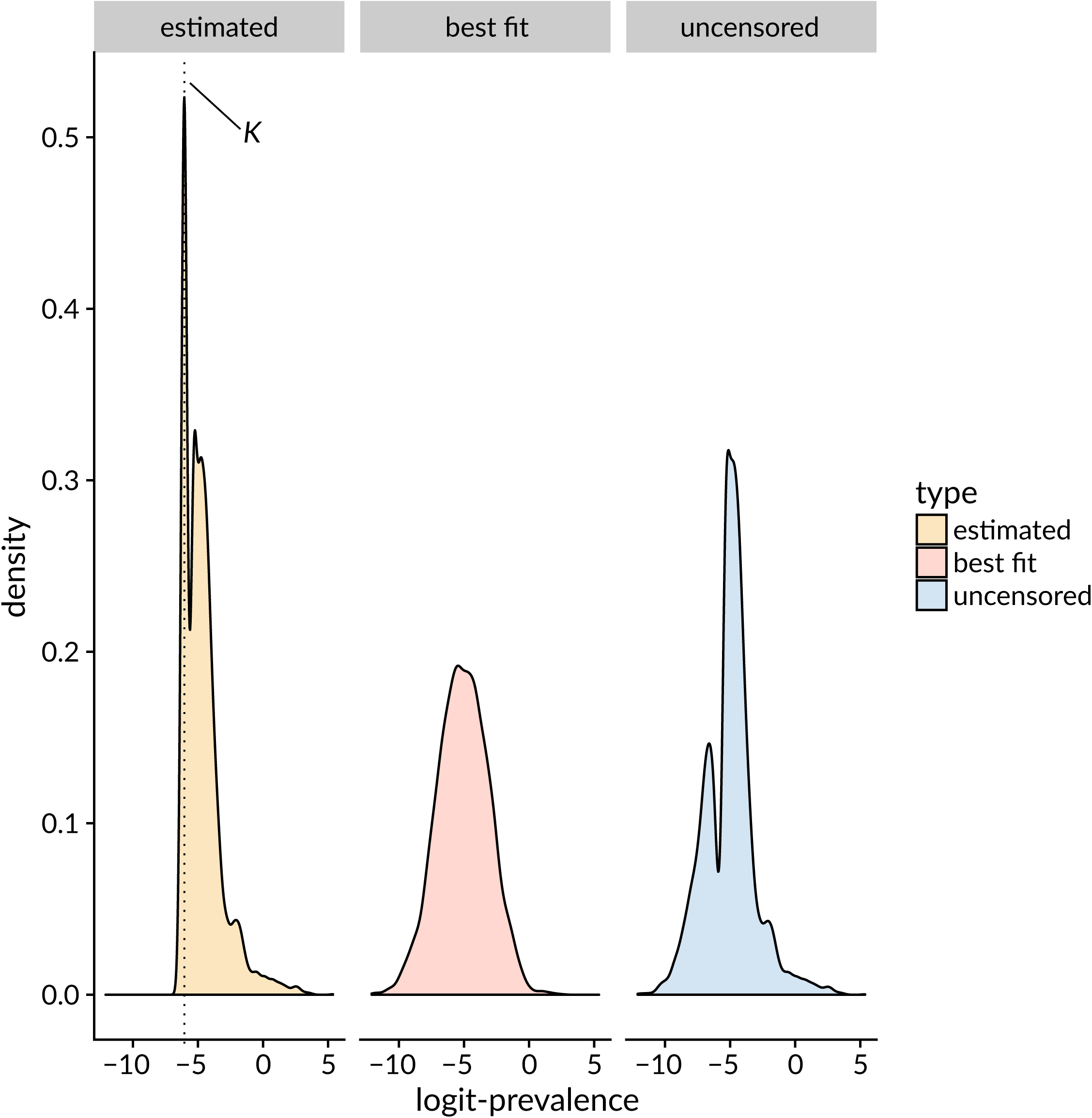
Distribution of estimated logit-prevalence 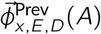, **showing impact of censoring.** A) Density of estimated logit-prevalence distribution, 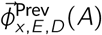, showing pile-up of values at the limit of detection *K*. B) Density of a normal distribution with mean and standard deviation obtained from best-fit of truncated normal to 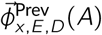. C) “Uncensored” version of 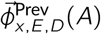. Data points at or below *K* have been replaced by random sampling from a truncated normal, with mean and standard deviation as in B and with *K* as upper truncation point.

**Figure S7.**
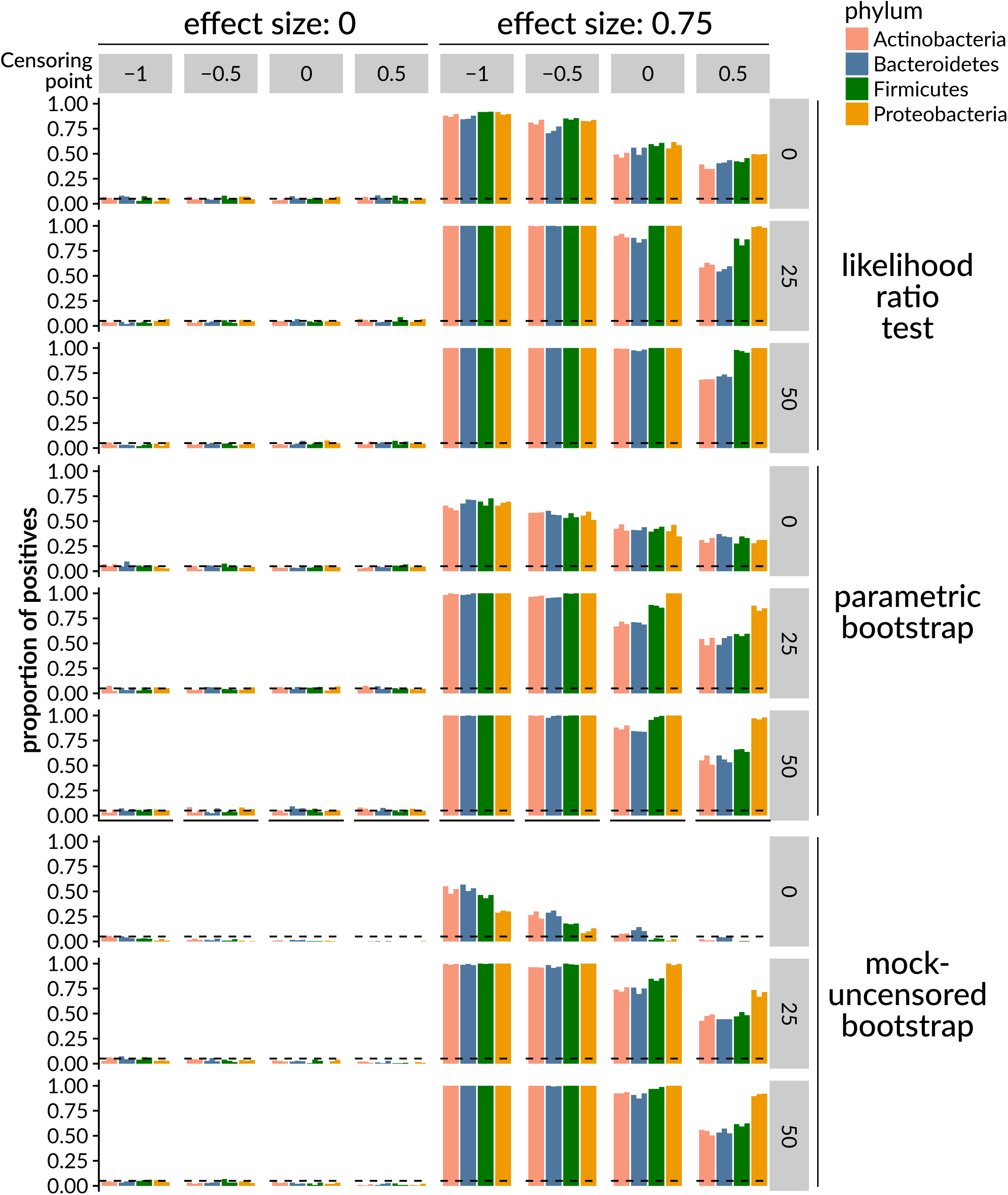
Impact of left-censoring on the false positive rate and power of phylogenetic tests. Bars give replicate measurements of false positive rate (left, effect size of 0) and power (right, effect size of 0.75) across the different phyla (colors), based on simulating binary genotypes and continuous phenotypes as in Methods 4.11, with varying levels of left-censoring (“censoring point”), and obtaining *p*-values with the three methods described in Methods 4.12. Horizontal dashed lines give a rate of 0.05. Binary genotypes had varying levels of Ives-Garland *α* (0, 25, 50), representing high to low phylogenetic signal.

**Figure S8.**
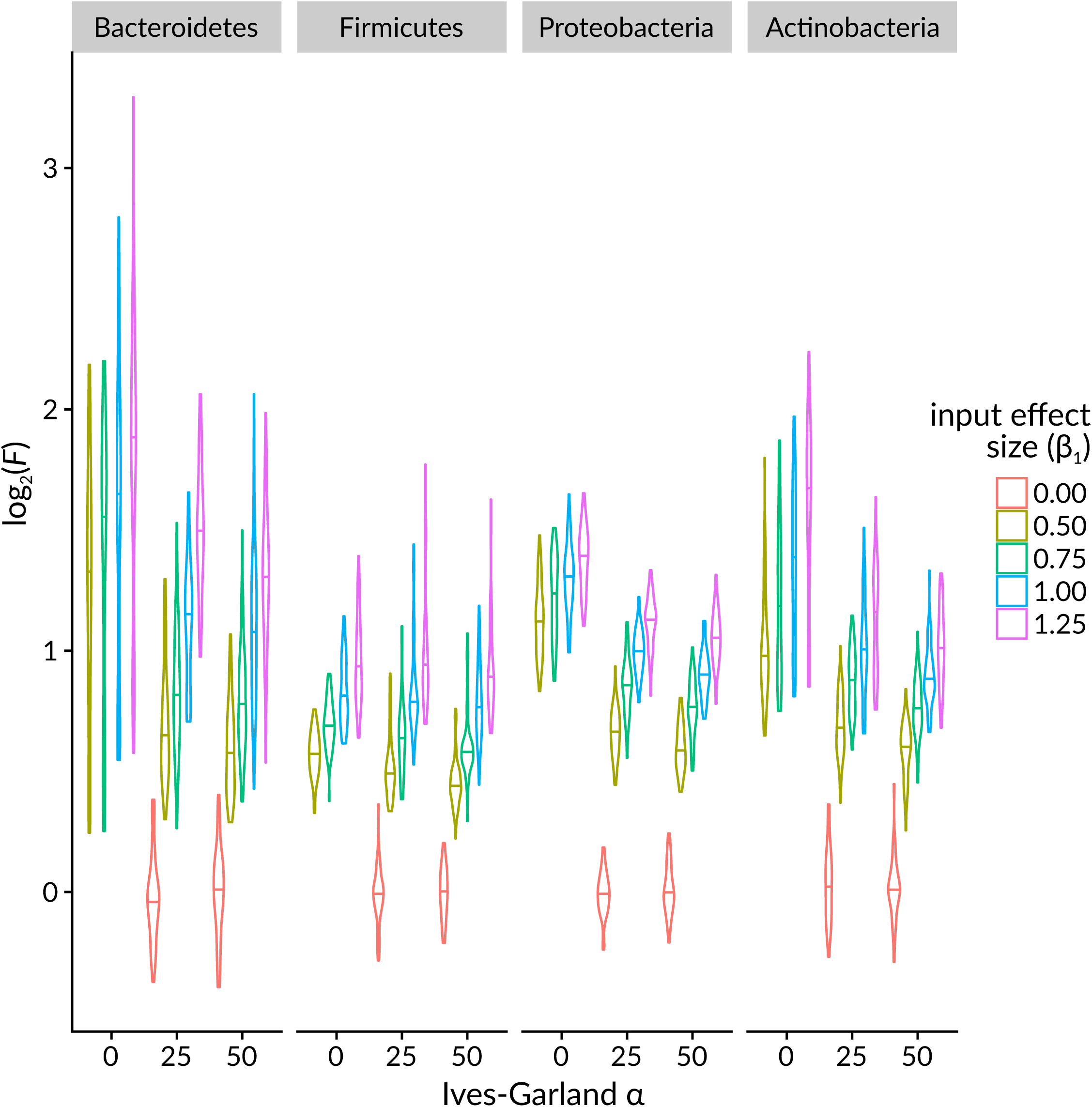
Simulations showing the prevalence ratios *F* corresponding to various effect sizes. To give a more intuitive sense of scale for simulated effect sizes, simulations were performed as in Methods 4.11, with effect sizes *β*_1_ ranging from 0 to After fitting phylogenetic models to the simulated phenotypes and genotypes, the average prevalences with the simulated gene, logistic(*β*_1,*g*_ + *β*_0,*g*_), and without, logistic(*β*_0,*g*_), were computed, and their ratio *F* was taken. log_2_(*F*) is plotted here, such that a value of 1 means the gene conferred (on average) a 2-fold change in prevalence. Violin plots were made of 50 simulations.

**Figure S9.**
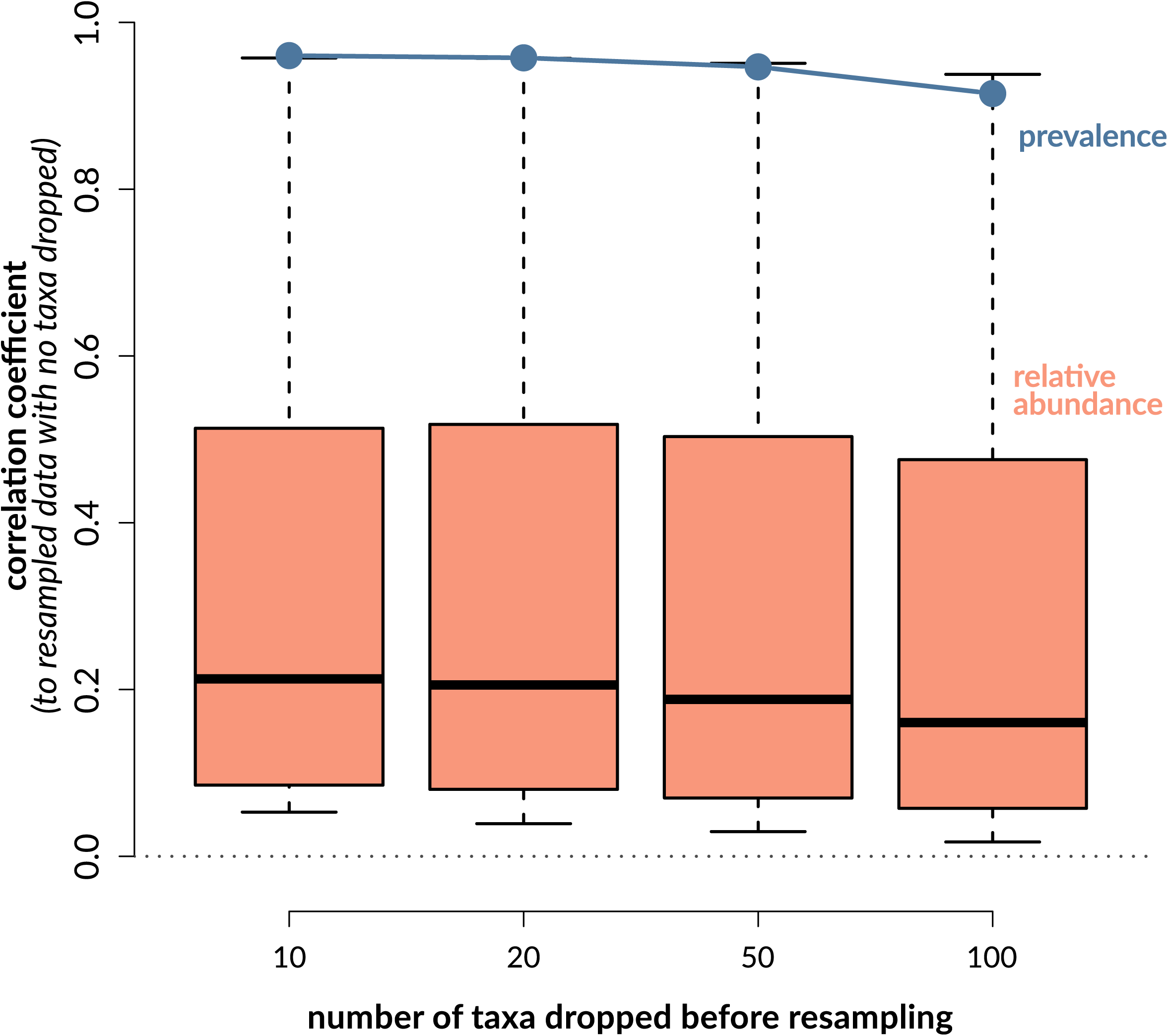
Effect of resampling after dropping abundant taxa on prevalence and abundance. Dirichlet-Multinomial sampling was performed on read counts either as-is or first dropping the 10, 20, 50, or 100 most-abundant taxa (X-axis). Relative abundances and prevalences were computed from all resampled datasets. The Y-axis represents Pearson’s correlation of prevalences (teal) or abundances (orange). Microbial prevalences with dropped taxa were compared to prevalence calculated without dropping taxa, yielding one correlation per sampling. For relative abundance profiles, the same microbe’s profile was compared between the as-is sampling and the sampling that dropped taxa, yielding a distribution of correlations (orange box-and-whisker plots). Each correlation coefficient is an average calculated from 100 resamplings.

## References

1. Slack E, Hapfelmeier S, Stecher B, Velykoredko Y, Stoel M, Lawson MAE, et al. Innate and adaptive immunity cooperate flexibly to maintain host-microbiota mutualism. Science. 2009;325(5940):617–620. doi:10.1126/science.1172747.

2. Atarashi K, Tanoue T, Shima T, Imaoka A, Kuwahara T, Momose Y, et al. Induction of colonic regulatory T cells by indigenous *Clostridium* species. Science. 2011;331(6015):337–341. doi:10.1126/science.1198469.

3. Mazmanian SK, Round JL, Kasper DL. A microbial symbiosis factor prevents intestinal inflammatory disease. Nature. 2008;453(7195):620–625. doi:10.1038/nature07008.

4. Sassone-Corsi M, Raffatellu M. No vacancy: how beneficial microbes cooperate with immunity to provide colonization resistance to pathogens. Journal of Immunology. 2015;194(9):4081–7. doi:10.4049/jimmunol.1403169.

5. Yano JM, Yu K, Donaldson GP, Shastri GG, Ann P, Ma L, et al. Indigenous bacteria from the gut microbiota regulate host serotonin biosynthesis. Cell. 2015;161(2):264–76. doi:10.1016/j.cell.2015.02.047.

6. Peng L, Li ZR, Green RS, Holzman IR, Lin J. Butyrate enhances the intestinal barrier by facilitating tight junction assembly via activation of AMP-activated protein kinase in Caco-2 cell monolayers. Journal of Nutrition. 2009;139(9):1619–1625. doi:10.3945/jn.109.104638.

7. Reber SO, Siebler PH, Donner NC, Morton JT, Smith DG, Kopelman JM, et al. Immunization with a heat-killed preparation of the environmental bacterium *Mycobacterium vaccae* promotes stress resilience in mice. Proceedings of the National Academy of Sciences. 2016;113(22):E3130–E3139. doi:10.1073/pnas.1600324113.

8. Garrett WS, Gallini CA, Yatsunenko T, Michaud M, DuBois A, Delaney ML, et al. *Enterobacteriaceae* act in concert with the gut microbiota to induce spontaneous and maternally transmitted colitis. Cell Host & Microbe. 2010;8(3):292–300. doi:10.1016/j.chom.2010.08.004.

9. Kullberg MC, Ward JM, Gorelick PL, Caspar P, Hieny S, Cheever A, et al. *Helicobacter hepaticus* triggers colitis in specific-pathogen-free interleukin-10 (IL-10)-deficient mice through an IL-12- and gamma interferon-dependent mechanism. Infection and Immunity. 1998;66(11):5157–66.

10. Kostic A, Chun E, Robertson L, Glickman J, Gallini C, Michaud M, et al. *Fusobacterium nucleatum* potentiates intestinal tumorigenesis and modulates the tumor-immune microenvironment. Cell Host & Microbe. 2013;14(2):207–215. doi:10.1016/j.chom.2013.07.007.

11. Bartlett JG. *Clostridium difficile*-associated enteric disease. Current Infectious Disease Reports. 2002;4(6):477–483.

12. van Nood E, Vrieze A, Nieuwdorp M, Fuentes S, Zoetendal EG, de Vos WM, et al. Duodenal infusion of donor feces for recurrent *Clostridium difficile*. New England Journal of Medicine. 2013;368(5):407–415. doi:10.1056/NEJMoa1205037.

13. Weingarden A, González A, Vázquez-Baeza Y, Weiss S, Humphry G, Berg-Lyons D, et al. Dynamic changes in short- and long-term bacterial composition following fecal microbiota transplantation for recurrent *Clostridium difficile* infection. Microbiome. 2015;3(1):10. doi:10.1186/s40168-015-0070-0.

14. Khanna S, Vazquez-Baeza Y, González A, Weiss S, Schmidt B, Muñiz-Pedrogo DA, et al. Changes in microbial ecology after fecal microbiota transplantation for recurrent *C. difficile* infection affected by underlying inflammatory bowel disease. Microbiome. 2017;5(1):55. doi:10.1186/s40168-017-0269-3.

15. Carvalho F, Koren O, Goodrich J, Johansson MV, Nalbantoglu I, Aitken J, et al. Transient inability to manage Proteobacteria promotes chronic gut inflammation in TLR5-deficient mice. Cell Host & Microbe. 2012;12(2):139–152. doi:10.1016/j.chom.2012.07.004.

16. Chassaing B, Koren O, Carvalho FA, Ley RE, Gewirtz AT. AIEC pathobiont instigates chronic colitis in susceptible hosts by altering microbiota composition. Gut. 2014;63(7):1069–1080. doi:10.1136/gutjnl-2013-304909.

17. Vilhjálmsson BJ, Nordborg M. The nature of confounding in genome-wide association studies. Nature Reviews Genetics. 2012;14(1):1–2. doi:10.1038/nrg3382.

18. Kim PJ, Price ND. Genetic co-occurrence network across sequenced microbes. PLoS Computational Biology. 2011;7(12):e1002340. doi:10.1371/journal.pcbi.1002340.

19. Porter SS, Chang PL, Conow CA, Dunham JP, Friesen ML. Association mapping reveals novel serpentine adaptation gene clusters in a population of symbiotic *Mesorhizobium*. The ISME Journal. 2017;11(1):248–262. doi:10.1038/ismej.2016.88.

20. Collins C, Didelot X. A phylogenetic method to perform genome-wide association studies in microbes that accounts for population structure and recombination. PLoS Computational Biology. 2018;14(2):e1005958. doi:10.1371/journal.pcbi.1005958.

21. Zhao N, Chen J, Carroll I, Ringel-Kulka T, Epstein M, Zhou H, et al. Testing in microbiome-profiling studies with MiRKAT, the microbiome regression-based kernel association test. The American Journal of Human Genetics. 2015;96(5):797–807. doi:10.1016/j.ajhg.2015.04.003.

22. Silverman JD, Washburne AD, Mukherjee S, David LA. A phylogenetic transform enhances analysis of compositional microbiota data. eLife. 2017;6. doi:10.7554/eLife.21887.

23. Tang ZZ, Chen G, Alekseyenko AV, Li H. A general framework for association analysis of microbial communities on a taxonomic tree. Bioinformatics. 2016;33(9):btw804. doi:10.1093/bioinformatics/btw804.

24. Lozupone CA, Hamady M, Cantarel BL, Coutinho PM, Henrissat B, Gordon JI, et al. The convergence of carbohydrate active gene repertoires in human gut microbes. Proceedings of the National Academy of Sciences. 2008;105(39):15076–15081. doi:10.1073/pnas.0807339105.

25. Felsenstein J. Phylogenies and the comparative method. The American Naturalist. 1985;.

26. Grafen A. The phylogenetic regression. Philosophical Transactions of the Royal Society of London Series B, Biological Sciences. 1989;326(1233):119–57.

27. Dunn CW, Zapata F, Munro C, Siebert S, Hejnol A. Pairwise comparisons across species are problematic when analyzing functional genomic data. bioRxiv. 2017;.

28. Levy A, Salas Gonzalez I, Mittelviefhaus M, Clingenpeel S, Herrera Paredes S, Miao J, et al. Genomic features of bacterial adaptation to plants. Nature Genetics. 2018;50(1):138–150. doi:10.1038/s41588-017-0012-9.

29. Ord TJ, Martins EP. Tracing the origins of signal diversity in anole lizards: phylogenetic approaches to inferring the evolution of complex behaviour. Animal Behaviour. 2006;71(6):1411–1429. doi:10.1016/j.anbehav.2005.12.003.

30. Zaneveld JRR, Parfrey LW, Van Treuren W, Lozupone C, Clemente JC, Knights D, et al. Combined phylogenetic and genomic approaches for the high-throughput study of microbial habitat adaptation. Trends in Microbiology. 2011;19(10):472–82. doi:10.1016/j.tim.2011.07.006.

31. Nayfach S, Rodriguez-Mueller B, Garud N, Pollard KS. An integrated metagenomics pipeline for strain profiling reveals novel patterns of bacterial transmission and biogeography. Genome Research. 2016;26(11):1612–1625. doi:10.1101/gr.201863.115.

32. Turnbaugh PJ, Hamady M, Yatsunenko T, Cantarel BL, Duncan A, Ley RE, et al. A core gut microbiome in obese and lean twins. Nature. 2009;457(7228):480–4. doi:10.1038/nature07540.

33. Human Microbiome Project Consortium. Structure, function and diversity of the healthy human microbiome. Nature. 2012;486(7402):207–14. doi:10.1038/nature11234.

34. Gevers D, Kugathasan S, Denson L, Vázquez-Baeza Y, Van Treuren W, Ren B, et al. The treatment-naive microbiome in new-onset Crohn’s disease. Cell Host & Microbe. 2014;15(3):382–392. doi:10.1016/j.chom.2014.02.005.

35. Meyer F, Overbeek R, Rodriguez A. FIGfams: yet another set of protein families. Nucleic Acids Research. 2009;37(20):6643–54. doi:10.1093/nar/gkp698.

36. Sakamoto M, Ohkuma M. Bacteroides reticulotermitis sp. nov., isolated from the gut of a subterranean termite (*Reticulitermes speratus*). International Journal of Systematic and Evolutionary Microbiology. 2013;63(Pt 2):691–695. doi:10.1099/ijs.0.040931-0.

37. Ives AR, Garland T. Phylogenetic logistic regression for binary dependent variables. Systematic Biology. 2010;59(1):9–26. doi:10.1093/sysbio/syp074.

38. Browne HP, Forster SC, Anonye BO, Kumar N, Neville BA, Stares MD, et al. Culturing of “unculturable” human microbiota reveals novel taxa and extensive sporulation. Nature. 2016;533(7604):543–546. doi:10.1038/nature17645.

39. Swick MC, Koehler TM, Driks A. Surviving between hosts: sporulation and transmission. Microbiology Spectrum. 2016;4(4). doi:10.1128/microbiolspec.VMBF-0029-2015.

40. De Biase D, Pennacchietti E. Glutamate decarboxylase-dependent acid resistance in orally acquired bacteria: function, distribution and biomedical implications of the *gadBC* operon. Molecular Microbiology. 2012;86(4):770–786. doi:10.1111/mmi.12020.

41. Srinivasa Rao PS, Lim TM, Leung KY. Functional genomics approach to the identification of virulence genes involved in *Edwardsiella tarda* pathogenesis. Infection and Immunity. 2003;71(3):1343–51.

42. Cotter PD, Gahan CG, Hill C. A glutamate decarboxylase system protects *Listeria monocytogenes* in gastric fluid. Molecular Microbiology. 2001;40(2):465–75.

43. Wargo MJ, Meadows JA. Carnitine in bacterial physiology and metabolism. Microbiology. 2015;161(6):1161–1174. doi:10.1099/mic.0.000080.

44. Staley C, Weingarden AR, Khoruts A, Sadowsky MJ. Interaction of gut microbiota with bile acid metabolism and its influence on disease states. Applied Microbiology and Biotechnology. 2017;101(1):47–64. doi:10.1007/s00253-016-8006-6.

45. Marques JC, Oh IK, Ly DC, Lamosa P, Ventura MR, Miller ST, et al. LsrF, a coenzyme A-dependent thiolase, catalyzes the terminal step in processing the quorum sensing signal autoinducer-2. Proceedings of the National Academy of Sciences. 2014;111(39):14235–14240. doi:10.1073/pnas.1408691111.

46. Thompson J, Oliveira R, Djukovic A, Ubeda C, Xavier K. Manipulation of the quorum sensing signal ai-2 affects the antibiotic-treated gut microbiota. Cell Reports. 2015;10(11):1861–1871. doi:10.1016/j.celrep.2015.02.049.

47. Wu M, McNulty NP, Rodionov DA, Khoroshkin MS, Griffin NW, Cheng J, et al. Genetic determinants of in vivo fitness and diet responsiveness in multiple human gut *Bacteroides*. Science. 2015;350(6256):aac5992–aac5992. doi:10.1126/science.aac5992.

48. Wang J, Jia H. Metagenome-wide association studies: fine-mining the microbiome. Nature Reviews Microbiology. 2016;14(8):508–522. doi:10.1038/nrmicro.2016.83.

49. Shin NR, Whon TW, Bae JW. Proteobacteria: microbial signature of dysbiosis in gut microbiota. Trends in Biotechnology. 2015;33(9):496–503. doi:10.1016/j.tibtech.2015.06.011.

50. Lynch SV, Pedersen O. the human intestinal microbiome in health and disease. The New England Journal of Medicine. 2016;375(24):2369–2379. doi:10.1056/NEJMra1600266.

51. Nielsen HB, Almeida M, Juncker AS, Rasmussen S, Li J, Sunagawa S, et al. Identification and assembly of genomes and genetic elements in complex metagenomic samples without using reference genomes. Nature Biotechnology. 2014;32(8):822–828. doi:10.1038/nbt.2939.

52. Li J, Jia H, Cai X, Zhong H, Feng Q, Sunagawa S, et al. An integrated catalog of reference genes in the human gut microbiome. Nature Biotechnology. 2014;32(8):834–841. doi:10.1038/nbt.2942.

53. Glasser AL, Boudeau J, Barnich N, Perruchot MH, Colombel JF, Darfeuille-Michaud A. Adherent invasive *Escherichia coli* strains from patients with Crohn’s disease survive and replicate within macrophages without inducing host cell death. Infection and Immunity. 2001;69(9):5529–37.

54. Barnich N, Boudeau J, Claret L, Darfeuille-Michaud A. Regulatory and functional co-operation of flagella and type 1 pili in adhesive and invasive abilities of AIEC strain LF82 isolated from a patient with Crohn’s disease. Molecular Microbiology. 2003;48(3):781–794. doi:10.1046/j.1365-2958.2003.03468.x.

55. Small CLN, Reid-Yu SA, McPhee JB, Coombes BK. Persistent infection with Crohn’s disease-associated adherent-invasive *Escherichia coli* leads to chronic inflammation and intestinal fibrosis. Nature Communications. 2013;4:1957. doi:10.1038/ncomms2957.

56. Lawley TD, Klimke WA, Gubbins MJ, Frost LS. F factor conjugation is a true type IV secretion system. FEMS Microbiology Letters. 2003;224(1):1–15.

57. Stecher B, Denzler R, Maier L, Bernet F, Sanders MJ, Pickard DJ, et al. Gut inflammation can boost horizontal gene transfer between pathogenic and commensal *Enterobacteriaceae*. Proceedings of the National Academy of Sciences. 2012;109(4):1269–1274. doi:10.1073/pnas.1113246109.

58. Ives AR, Helmus MR. Generalized linear mixed models for phylogenetic analyses of community structure. Ecological Monographs. 2011;81(3):511–525. doi:10.1890/10-1264.1.

59. Konstantinidis KT, Tiedje JM. Genomic insights that advance the species definition for prokaryotes. Proceedings of the National Academy of Sciences. 2005;102(7):2567–72. doi:10.1073/pnas.0409727102.

60. Jain C, Rodriguez-R LM, Phillippy AM, Konstantinidis KT, Aluru S. High-throughput ANI analysis of 90K prokaryotic genomes reveals clear species boundaries. bioRxiv. 2017; p. 225342. doi:10.1101/225342.

61. Rosselló-Móra R, Amann R. Past and future species definitions for Bacteria and Archaea. Systematic and Applied Microbiology. 2015;38(4):209–216. doi:10.1016/j.syapm.2015.02.001.

62. Cadillo-Quiroz H, Didelot X, Held NL, Herrera A, Darling A, Reno ML, et al. Patterns of gene flow define species of thermophilic archaea. PLoS Biology. 2012;10(2):e1001265. doi:10.1371/journal.pbio.1001265.

63. Richter M, Rosselló-Móra R. Shifting the genomic gold standard for the prokaryotic species definition. Proceedings of the National Academy of Sciences. 2009;106(45):19126–31. doi:10.1073/pnas.0906412106.

64. Winter DJ. rentrez: An R package for the NCBI eUtils API. PeerJ Preprints. 2017;doi:10.7287/peerj.preprints.3179v2.

65. Edgar RC. Search and clustering orders of magnitude faster than BLAST. Bioinformatics. 2010;26(19):2460–2461. doi:10.1093/bioinformatics/btq461.

66. Wattam AR, Abraham D, Dalay O, Disz TL, Driscoll T, Gabbard JL, et al. PATRIC, the bacterial bioinformatics database and analysis resource. Nucleic Acids Research. 2014;42(Database issue):D581–91. doi:10.1093/nar/gkt1099.

67. Price MN, Dehal PS, Arkin AP. FastTree 2–approximately maximum-likelihood trees for large alignments. PLoS ONE. 2010;5(3):e9490. doi:10.1371/journal.pone.0009490.

68. Edgar RC. MUSCLE: multiple sequence alignment with high accuracy and high throughput. Nucleic Acids Research. 2004;32(5):1792–1797. doi:10.1093/nar/gkh340.

69. Paradis E, Claude J, Strimmer K. APE: Analyses of Phylogenetics and Evolution in R language. Bioinformatics. 2004;20(2):289–290. doi:10.1093/bioinformatics/btg412.

70. Garamszegi LZ, Gonzalez-Voyer A. Working with the Tree of Life in Comparative Studies: How to Build and Tailor Phylogenies to Interspecific Datasets. In: Modern phylogenetic comparative methods and their application in evolutionary biology. Berlin, Heidelberg: Springer Berlin Heidelberg; 2014. p. 19–48. Available from: http://link.springer.com/10.1007/978-3-662-43550-2{_}2.

71. Karlsson FH, Tremaroli V, Nookaew I, Bergström G, Behre CJ, Fagerberg B, et al. Gut metagenome in European women with normal, impaired and diabetic glucose control. Nature. 2013;498(7452):99–103. doi:10.1038/nature12198.

72. Qin J, Li Y, Cai Z, Li S, Zhu J, Zhang F, et al. A metagenome-wide association study of gut microbiota in type 2 diabetes. Nature. 2012;490(7418):55–60. doi:10.1038/nature11450.

73. Zhu Y, Stephens RM, Meltzer PS, Davis SR. SRAdb: query and use public next-generation sequencing data from within R. BMC Bioinformatics. 2013;14(1):19. doi:10.1186/1471-2105-14-19.

74. Leinonen R, Akhtar R, Birney E, Bower L, Cerdeno-Tarraga A, Cheng Y, et al. The European Nucleotide Archive. Nucleic Acids Research. 2011;39(Database):D28–D31. doi:10.1093/nar/gkq967.

75. si Tung Ho L, Ané C. A linear-time algorithm for gaussian and non-gaussian trait evolution models. Systematic Biology. 2014;63(3):397–408. doi:10.1093/sysbio/syu005.

76. Storey JD, Bass AJ, Dabney A, Robinson D. qvalue: Q-value estimation for false discovery rate control; 2015. Available from: http://github.com/jdstorey/qvalue.

77. Storey JD, Tibshirani R. Statistical significance for genomewide studies. Proceedings of the National Academy of Sciences. 2003;100(16):9440–5. doi:10.1073/pnas.1530509100.

78. Kappelman MD, Rifas-Shiman SL, Kleinman K, Ollendorf D, Bousvaros A, Grand RJ, et al. The prevalence and geographic distribution of Crohn’s disease and ulcerative colitis in the United States. Clinical Gastroenterology and Hepatology. 2007;5(12):1424–1429. doi:10.1016/j.cgh.2007.07.012.

79. Aitchison J. The statistical analysis of compositional data. 1986;.

80. Overbeek R, Begley T, Butler RM, Choudhuri JV, Chuang HY, Cohoon M, et al. The subsystems approach to genome annotation and its use in the project to annotate 1000 genomes. Nucleic Acids Research. 2005;33(17):5691–5702. doi:10.1093/nar/gki866.

81. Hochberg Y, Benjamini Y. Controlling the false discovery rate: a practical and powerful approach to multiple testing. Journal of the Royal Statistical Society Series B (Methodological). 1995;1:289–300.

82. Subramanian A, Tamayo P, Mootha VK, Mukherjee S, Ebert BL, Gillette MA, et al. Gene set enrichment analysis: a knowledge-based approach for interpreting genome-wide expression profiles. Proceedings of the National Academy of Sciences of the United States of America. 2005;102(43):15545–50. doi:10.1073/pnas.0506580102.

